# Amphipathic helices sense the inner nuclear membrane environment through lipid packing defects

**DOI:** 10.1101/2024.11.14.623600

**Authors:** Shoken Lee, Anabel-Lise Le Roux, Mira Mors, Stefano Vanni, Pere Roca-Cusachs, Shirin Bahmanyar

## Abstract

Amphipathic helices (AHs) are ubiquitous protein motifs that modulate targeting to organellar membranes by sensing differences in bulk membrane properties. However, the adaptation between membrane-targeting AHs and the nuclear membrane environment that surrounds the genome is poorly understood. Here, we computationally screened for candidate AHs in a curated list of characterized and putative human inner nuclear membrane (INM) proteins. Cell biological and *in vitro* experimental assays combined with computational calculations demonstrated that AHs detect lipid packing defects over electrostatics to bind to the INM, indicating that the INM is loosely packed under basal conditions. Membrane tension resulting from hypotonic shock further promoted AH binding to the INM, whereas cell-substrate stretch did not enhance recruitment of membrane tension-sensitive AHs. Together, our work demonstrates the rules driving lipid-protein interactions at the INM, and its implications in the response of the nucleus to different stimuli.

## Introduction

Membrane-bound organelles have characteristic biophysical properties driven by their distinct lipid compositions. The endoplasmic reticulum (ER) and Golgi apparatus are defined by loose lipid packing and low electrostatics, whereas organelles of the late secretory pathway, plasma membrane, and mitochondria generally contain more tightly packed lipids and are rich with negatively charged lipids (Bigay and Antonny, 2012; Ernst et al., 2016; Holthuis and Menon, 2014). Protein regions called amphipathic helices (AHs) recognize these differences in bulk membrane properties and insert in the polar-apolar region of a single leaflet of a lipid bilayer (Drin and Antonny, 2010; Giménez-Andrés et al., 2018). The composition of the hydrophilic and hydrophobic amino acid residues that make up each face of the helix determines whether an AH detects membrane electrostatics or loose lipid packing (Drin et al., 2007; Hofbauer et al., 2018; Pranke et al., 2011) – this mechanism is distinct from the recognition of specific lipid species based on the head group composition (Lemmon, 2008). Loose lipid packing results from an increased level of unsaturated fatty acids or cone-shaped lipids, which induces lipid packing defects – regions in which hydrophobic tails are exposed to the aqueous cytoplasm (Bigay and Antonny, 2012; Holthuis and Menon, 2014). AHs with few charged amino acids on their hydrophilic face bind membranes through the highly sensitive detection of membrane packing defects driven by hydrophobic interactions (Drin et al., 2007; Pranke et al., 2011; Vanni et al., 2014). Some AHs (e.g., Amphipathic Lipid-Packing Sensor (ALPS) motifs) with no to very few charged amino acids are exquisite sensors of membrane curvature, the geometry of which intrinsically contains a high density of packing defects (Drin et al., 2007; Pranke et al., 2011; Vanni et al., 2014).

While a lot is known about how AHs detect differences in bulk membrane properties, little is known about whether these detection modes are relevant to the nuclear envelope (NE). The NE is unique among organelles because it is a relatively flat, uniform membrane sheet, with the exception of curved membranes at fusion points between the inner and outer nuclear membrane, which are highly occupied by nuclear pore complex proteins (Bahmanyar and Schlieker, 2020; Cornell and Antonny, 2018), and at relatively infrequent invaginations of the NE (e.g., nucleoplasmic reticulum (McPhee et al., 2024)). In addition, the NE shares a single membrane with the ER, which makes isolating the NE biochemically challenging and hinders direct determination of its lipid composition. The recent development of fluorescent INM lipid biosensors bypassed this challenge and indicated that the NE across multiple organisms contains the cone-shaped lipid diacylglycerol (DAG) (Foo et al., 2023; Lee et al., 2023; Romanauska and Köhler, 2018). Thus, the nuclear membrane may be loosely packed like the ER despite its flat structure. A lipid biosensor for phosphatidylserine (PS) suggested that the surface of the INM may also be defined by electrostatics (Niu et al., 2024). The degree to which negative charge versus packing defects define the membrane properties of the NE remains an important issue to resolve.

AHs are found in many nucleoporins, consistent with the highly curved feature of the nuclear pore membrane (Amm et al., 2023; Hamed and Antonin, 2021; Kralt et al., 2022). Less is known about how membrane-binding AHs detect the relatively flat surface of the INM. The best-known example of AH detection of the INM at non-NPC regions is by the soluble, rate-limiting enzyme in phosphocholine synthesis, CCTa/PCYT1A (Cornell and Antonny, 2018; Cornell, 2016; Haider et al., 2018). The resident INM nucleo-cytoskeletal protein Sun2 was also recently shown to contain an AH with a preference for packing defects (Lee et al., 2023). These examples suggest that the detection of bulk INM properties by AHs may be a more pervasive mechanism linked to NE-dependent processes.

Defining bulk INM properties and how they are sensed is particularly important when it comes to mechanical forces imposed on the NE. Recently it was shown that NE tension resulting from hypotonic shock or from physical confinement promotes INM-association of the C2 domain of the enzyme cPLA2 (Enyedi et al., 2016; Lomakin et al., 2020; Venturini et al., 2020). It has been proposed that cPLA2 recognizes increased packing defects at the INM; however, the extent to which different mechanical inputs affect lipid order at the INM beyond this example remains unexplored.

Prior screening approaches have revealed generalized characteristics of a subset of AHs and how they recognize a specific property of the target membrane (Delic et al., 2024; Drin et al., 2007; Prévost et al., 2018; van Hilten et al., 2023; van Hilten et al., 2022). Here, we identified previously unknown proteins by screening putative and known transmembrane-containing NE-associated proteins for membrane-sensing AH motifs by taking advantage of a deep-learning algorithm called MemBrain (Feng et al., 2022). We defined determinants in these AHs that facilitate binding to the INM and used multidisciplinary approaches to demonstrate that they preferentially associate with membrane packing defects. We utilized the property of AHs as detectors of lipid membrane properties to determine the extent of induction of INM lipid packing defects in response to distinct mechanical inputs. Our findings reveal that the INM is an unpacked membrane territory recognized by AHs, and demonstrates for the first time how different mechanical stimuli effect the detection of lipid packing defects at the NE.

## Results

Our prior search used machine-learning-based algorithms to yield seven predicted AHs in characterized inner nuclear membrane proteins, including the LEM-domain family, LINC complex proteins, Lamin B Receptor and NEMP1 (Lee et al., 2023). We set out to curate a larger number of proteins shown to associate with the NE to gain a better understanding of the amino acid-sequence code used by AHs to detect the flat membrane territory of the NE. To generate a more comprehensive list of known and previously uncharacterized nuclear membrane-associated proteins, we compiled data from biochemical studies in which the NE proteome was determined in distinct tissue types (Cheng et al., 2019; Korfali et al., 2010; Korfali et al., 2012; Schirmer et al., 2003; Wilkie et al., 2011) and obtained a list of 410 proteins (Fig. 1A; see Fig. S1A and S1B and Methods for methodological details and Tables S1-S5 for the complete lists of the proteins). We refined this list to 281 proteins by prioritizing those in which more than one publication supported NE-association or additional support for NE-association was available from the Human Protein Atlas or UniProt (Fig. S1A and Methods; see Tables S6-S7 for the complete lists of proteins). We leveraged an *in-silico* deep-learning-based algorithm called MemBrain to predict AHs (Feng et al., 2022) within our extensive list of proteins. We filtered their published predictions that covered 11,750 putative transmembrane proteins from various organisms, hereafter referred to as the “MemBrain list,” to only those from humans (Fig. S1A and Tables S8-S9). Merging the compiled list of nuclear membrane proteins with the filtered MemBrain analysis resulted in 87 proteins that contained either one or two predicted AHs (Fig. 1; Fig. S1A; Tables S10-S11).

**Fig. 1.**
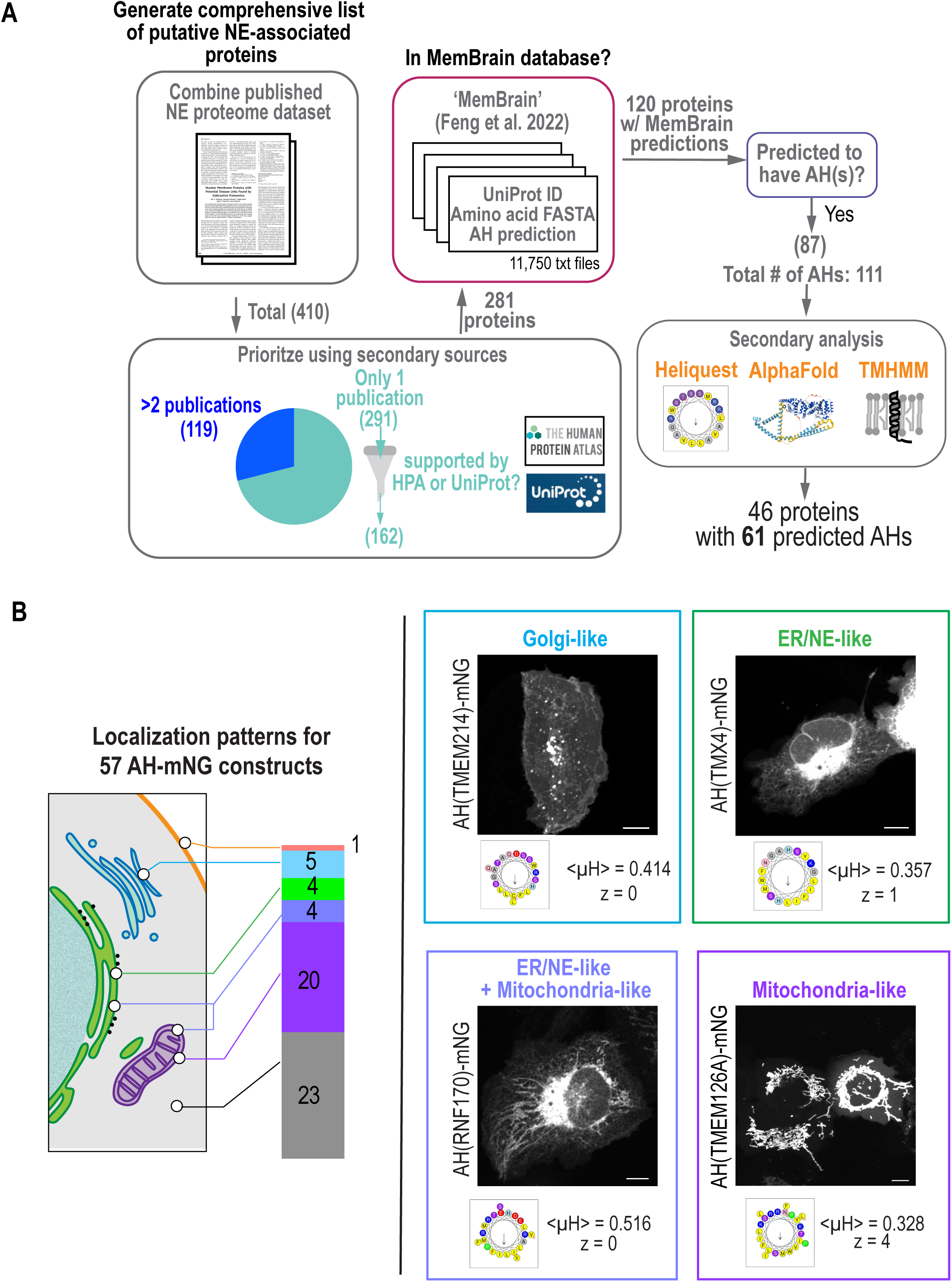
Patterns of localization of candidate AHs identified in curated list of putative nuclear envelope-associated proteins. (A) Flow chart of data mining for putative nuclear envelope proteins and identification of predicted amphipathic helices (AHs). See Fig. S1 for more detail. (B) Breakdown of AH-mNG localization patterns with corresponding color-coded schematic. Spinning disk confocal images of U2OS cells expressing indicated constructs represent example localization patterns. Wheel projections of the predicted AHs with mean hydrophobic moment <µH> and net charge z is shown. See Fig. S3, S5 and S6 for representative images of other AH-mNG candidates. Scale bars, 10 µm.

Similar to our prior analyses of AHs in characterized NE proteins (Lee et al., 2023), AHs were identified from the ‘MemBrain’ list in Sun2, Sun1, NEMP1, LEMD2, and Man1 (Fig. S1C). There was no overlap between the lists of AH candidates from our study and the ALPS-like motifs identified in the proteome-wide bioinformatics screen from Drin et al. 2007 (Fig. S2) – ‘MemBrain’ analysis was not available for the only four NE proteins that overlapped between the two screens for which they identified to contain candidate ALPS-like motifs (Fig. S2). We eliminated potential false positives in our list to confirm that putative AH regions are predicted helices by AlphaFold2 (Jumper et al., 2021), to eliminate regions predicted as transmembrane regions by TMHMM (Krogh et al., 2001), and to confirm their strong amphiphilicity as predicted by HeliQuest (Gautier et al., 2008). Our final list included 46 proteins with 61 promising AH candidates (Fig. 1A and Table S12).

We next tested whether our list of putative AHs associate with cytosolic membrane-bound organelles that have well-defined lipid territories. We reasoned that an understanding of whether AHs from NE-associated proteins bind different lipid territories of cytosolic membrane organelles could inform our understanding of the properties of the nuclear membrane. We successfully cloned 57 AH candidates appended to mNeonGreen (mNG) fluorescent protein at their C-termini and determined their localizations when expressed in living U2OS cells. Twenty-three AHs localized to the cytosol and nucleoplasm, similar to mNG alone (Fig. 1B; Fig. S3), and this was not because the mNG was abnormally cleaved from the AH sequence (Fig. S4). The 34 AHs that localized to membranes displayed varied localization patterns that resembled mitochondria, ER, and Golgi, and one AH localized to the cell periphery resembling the plasma membrane (Fig. 1B; Figs. S5-S7; Table S13). AH-mNG proteins are small enough to diffuse into the nucleus; however, none of the AHs exclusively localized to the INM. This is in line with prior work demonstrating that full-length proteins direct the organellar localization of AHs, whereas expressing AHs on their own reveals their independent membrane binding preferences (Doucet et al., 2015; Lee et al., 2023; Pranke et al., 2011; Prévost et al., 2018). Note that a limitation of our approach is that an N-terminal AH could potentially serve as a mitochondrial import signal (Abe et al., 2000).

We next determined if there was a pattern in the amino acid sequences of AHs that could suggest an adaptation to different lipid territories. There were no differences in length, net charge, or hydrophobic moment that clearly discriminated AHs (Fig. 2A-2C). We also compared the “D factor” for AH sequences, which serves as a prediction tool for the likelihood of association to lipid bilayers through a combined hydrophobic moment and net charge calculation (possible association if D > 0.65 and likely association if D > 1.3) (Gautier et al., 2008), but did not find any major differences between the different groupings (Fig. 2D).

**Fig. 2.**
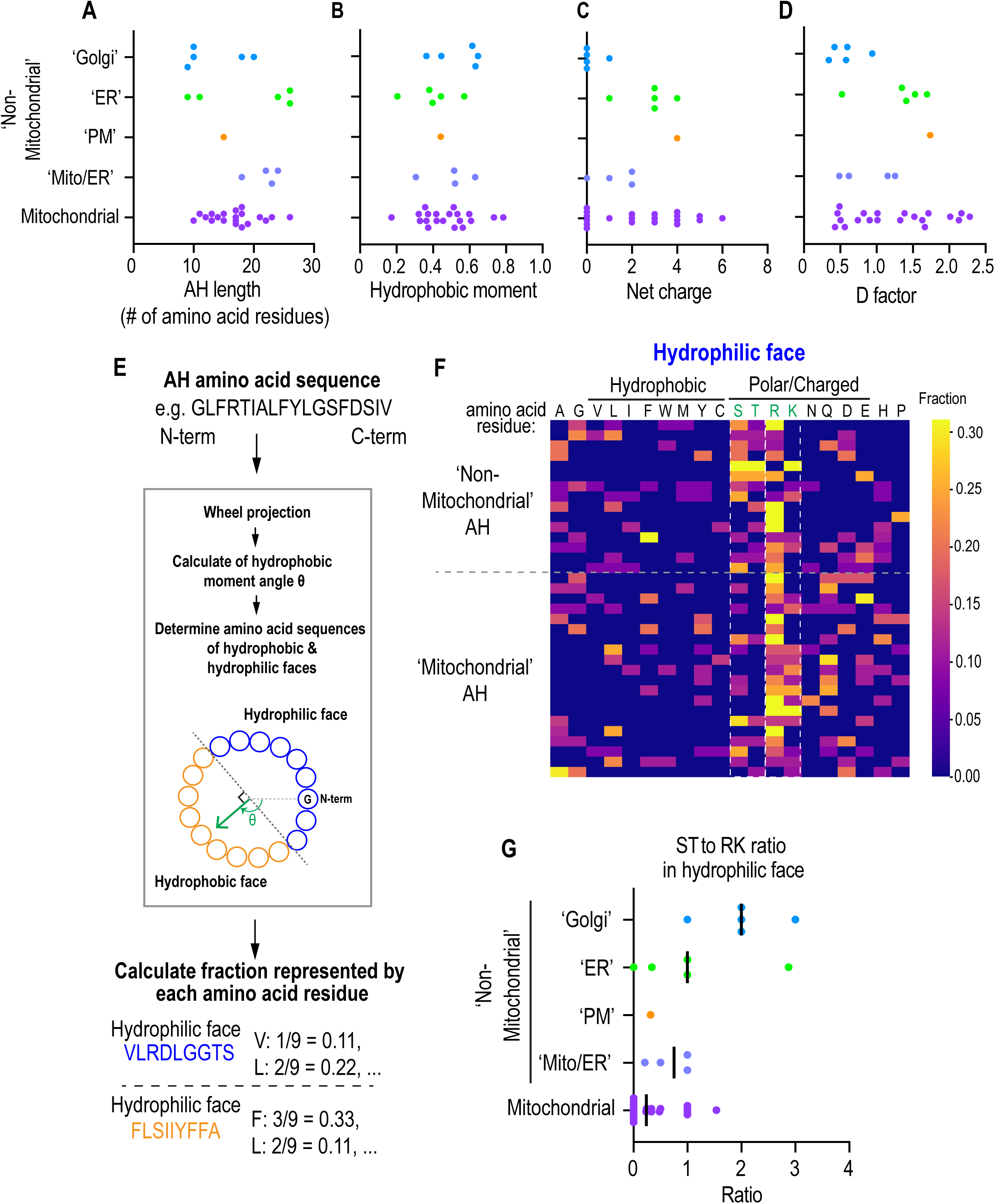
Analysis of amino acid sequences of predicted AHs categorized by their localization patterns. (A-D) Scatter plot of the indicated metrices of AH candidates categorized by their localization patterns as determined from experiments represented in Fig. 1B. (E) Flow chart for determining the hydrophilic or hydrophobic face of each AH and the calculation of the fraction of each amino acid present within either face. (F) Heat map representation of the fraction of each amino acid (columns) present within the hydrophilic face of each amphipathic helix (row) categorized by indicated localization patterns. Outlined are S/T and R/K residues. (G) Scatter plot representing the ratio of S/T residues to R/K residues in the hydrophilic face of each amphipathic helix categorized by localization as determined from experiments represented in Fig. 1B. Bar represents median value.

A trend was revealed from analysis of the amino acid compositions of the hydrophilic and hydrophobic faces of 32 AHs (Fig. 2E and 2F; Fig. S8) in which ER, Golgi or plasma membrane (‘non-mitochondrial’)-localized AHs were enriched in Ser/Thr residues in the hydrophilic face (Fig. 2F) and enriched in Leu residue relative to Ile/Phe/Trp residues in the hydrophobic face (Fig. S8). Ser/Thr residues in the hydrophilic face of AHs in place of other charged residues, such as Arg/Lys, suggests that the driving force for membrane binding is through the hydrophobic effect rather than electrostatic charges (Bigay and Antonny, 2012). This is exemplified in the ALPS motifs of ArfGAP1 and GMAP210 that contain few Arg/Lys residues in the hydrophilic faces – the prerequisite for membrane binding for ALPS motifs is lipid packing defects, making them exquisite sensors of membrane curvature (Drin et al., 2007; Pranke et al., 2011; Vanni et al., 2014; Vanni et al., 2013). Indeed, Golgi and ER-localized AHs contained a higher ratio of Ser/Thr residues to Arg/Lys residues in the hydrophilic face, indicating a preference for membrane territories with lipid packing defects (Fig. 2G).

Directing ten of the AH-mNG constructs to the nucleus by appending a nuclear localization signal (NLS) to their C-termini revealed INM localization, distinct from the mNG-NLS control, for AHs that localize to the Golgi, ER or plasma membrane when expressed in the cytosol (Fig. 3A-3C). The stronger enrichment at the INM was confirmed with semiautomated image analysis that determines the ratio of fluorescent signal at the nuclear rim and inside the nucleus (‘NE enrichment score’) (Fig. 3D; see Methods). AH-mNG constructs appended to an NLS that localize to mitochondria when expressed in the cytosol did not associate with the nuclear rim – when directed to the nucleus, these protein motifs localized to the nucleoplasm and nuclear bodies in a manner that resembled the distribution of the mNG-NLS control (Fig. 3B-3D). These results suggest that electrostatics alone are not sufficient to promote INM binding because ‘mitochondrial’ AH-mNG constructs contain low S/T:R/K ratios in their hydrophilic face (Fig. 2F and 2G). Interestingly, the ALPS motif of Nup133 (Drin et al., 2007; Nordeen et al., 2020) that has a high S/T:R/K ratio as well as the ALPS motifs of ArfGAP1 (Bigay et al., 2005; Mesmin et al., 2007) that also has few charged residues were less associated with the INM relative to the AH of Nup153 (Vollmer et al., 2015) and the other INM-associated AHs tested (Fig. 3; Fig. S9). Thus, a combination of electrostatics and membrane packing defects is likely required for robust AH association with the INM under basal conditions.

**Fig. 3.**
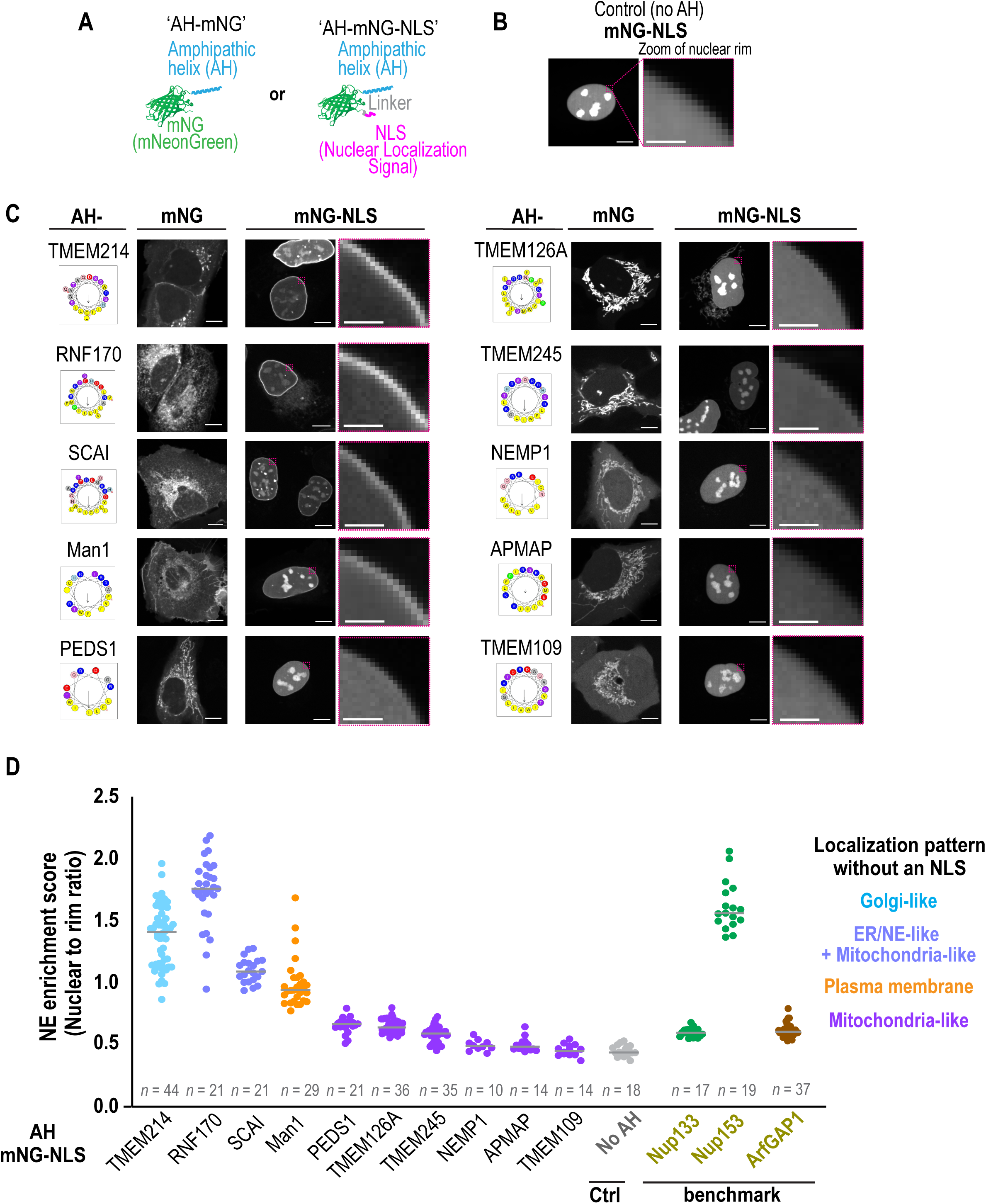
Appending candidate AHs with a nuclear localization signal (NLS) reveals their propensity to associate with the inner nuclear membrane. (A) Schematic representation of AH-mNG and AH-mNG-NLS protein design. (B-C) Representative spinning disk confocal images of a single section in living U2OS cells expressing mNG-NLS alone (in B) or the indicated AH-mNG and corresponding AH-mNG-NLS protein in (C) with magnified zoom of nuclear rim area shown. Schematic representations of corresponding helical wheel projections and hydrophobic moment of candidate AH sequences are shown. (D) NE enrichment score of indicated AH-mNG-NLS proteins quantified from single z-slice of spinning disk confocal images. Categorization is based on non-NLS appended localization patterns of each candidate AH. NE enrichment scores of AH sequences from Nup133, Nup153, and ArfGAP1 are also shown as benchmark values (see Fig. S9, Methods, and Fig. S13). Data points from two or three independent experiments are shown. n = number of nuclei. Bars indicate median values. Scale bars, 10 µm or 2 µm (zoom).

We directly tested the role that electrostatics and lipid packing defects play in promoting AH binding to the INM by increasing the S/T:R/K ratio in a non-INM localized AH. Mutating R/K amino acids in the hydrophilic face of the AH of TMEM126 to S/T residues incrementally increased its INM association in cells (Fig. 4A). Wild-type and K11S/R15S mutated AH peptides tagged with monomeric GFP were purified from bacterial cells (Fig. S10) and their interactions with giant unilamellar vesicles (GUVs), which have very low intrinsic local curvature similar to the INM, composed of different lipid compositions were analyzed. We prepared fluorescently-labeled GUVs with lipid compositions that represent low packing defects (dioleoyl-phosphatidylcholine (DOPC) (‘PC-alone’)); a highly charged lipid bilayer (10% dioleoyl-phosphatidylserine (DOPS) (‘+PS’) or 10% dioleoyl-phosphatidylinositol (DOPI) (‘+PI’)); and a lipid bilayer with packing defects (10% dioleoyl-glycerol (DOG) (‘+DAG’) (Vamparys et al., 2013; Vanni et al., 2014) or saturated, methyl-branched chains phospholipid: 10-100 % diphytanoyl-phosphatidylcholine (‘+Branched PC’) (Čopič et al., 2018; Garten et al., 2015)). Neither the wild-type nor the K11S/R15S mutant AH bound to PC-alone GUVs, while both the wild-type and mutant AH bound to the +PS or +PI GUVs, with the wild-type displaying greater association (Fig. 4B). These results are in line with electrostatics playing a role in membrane association through the positively charged amino acids in the hydrophilic face of both constructs (Drin et al., 2007; Pranke et al., 2011). Hydrophobic driving forces overcame the need for electrostatics only for K11S/R15S mutated AH peptides, which robustly associated with GUVs containing either the cone-shaped lipid DAG or branched PC known to induce packing defects (Fig. 4B and 4C) (Čopič et al., 2018; Garten et al., 2015; Vamparys et al., 2013; Vanni et al., 2014). The fact that only the K11S/R15S mutant, and not the wild-type AH, bound to GUVs containing packing defects suggests that a higher S/T:R/K ratio in the hydrophilic face of the AH of TMEM126 promoted membrane binding when nuclear-localized because the INM contains packing defects.

**Fig. 4.**
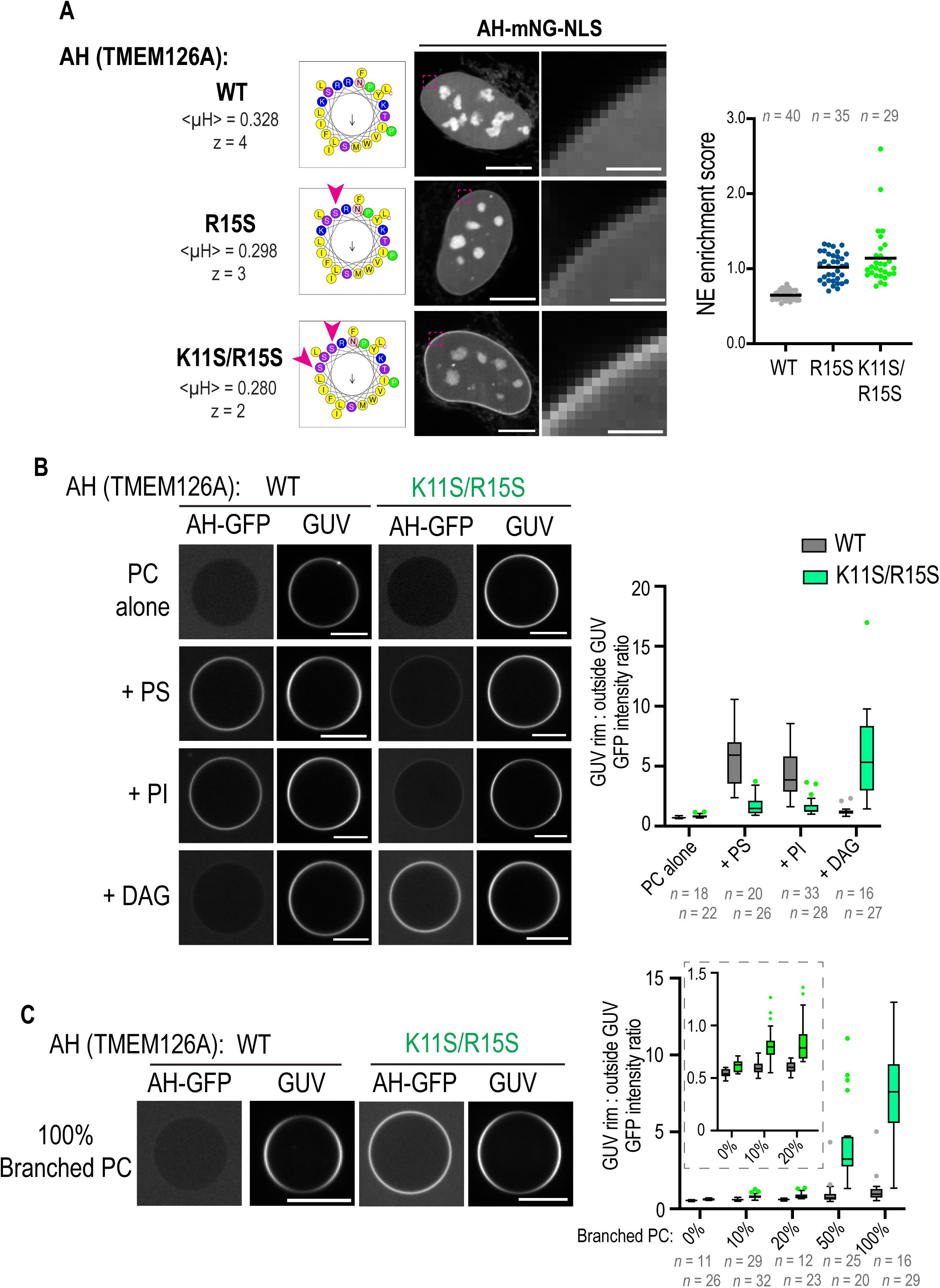
A higher ratio of S/T to R/K amino acid residues in the hydrophilic face of a candidate AH promotes its sensitivity to packing defects and association to the inner nuclear membrane. (A) Spinning disk confocal images of mNG channel in living U2OS cells expressing indicated wild type and mutated versions of AH(TMEM126)-mNG-NLS. Scatter plot (right) represents NE enrichment scores for corresponding AH sequences. n = number of nuclei. Data are pooled from two independent experiments. Scale bars, 10 µm or 2 µm (zoom). (B and C) Spinning disk confocal images of single z-slice of GUVs incubated with indicated AH-GFP recombinant protein. Box plot represents the ratio of fluorescence intensities of AH-GFP localization with GUV rim to outside GUV (see Methods and Fig. S13). In (C), values for branched PC 0-20% plots are magnified. n = number of GUVs. Data are pooled from two or three independent experiments. Scale bars, 10 µm.

To further support the idea that the presence of packing defects contributes to the driving force for INM-association of AHs, we used PMIpred, a computational predictor based on a transformer neural network trained on data derived from molecular dynamics simulations (van Hilten et al., 2024). PMIpred calculates the difference in free energy reduction upon AH helix association to ‘stretched’ and ‘non-stretched’ membrane bilayers containing a constant number of lipids (*ΔΔF*) (Fig. 5A) (van Hilten et al., 2023; van Hilten et al., 2022; van Hilten et al., 2024) based solely on peptide sequence. Since ‘stretched’ membranes result in the exposure of acyl chains to the solvent and to the increase of packing defects (Pinot et al., 2018; van Hilten et al., 2020), high *ΔΔF* values indicate that hydrophobicity is the main driving force for membrane association and hence packing-defect sensing. We compared the *ΔΔF* for our AH peptide candidates (57 in total tested by live imaging (Fig. 1)) to those of characterized AHs (van Hilten et al., 2023), which fall into three categories defined by their sensitivity to packing defects: binders (any membrane independent of curvature-induced packing defects), sensors (only bind curved membranes) and non-binders (don’t bind membranes) (van Hilten et al., 2023). We found that the AHs in our list that localized to cytosolic organelle membranes (34 in total) tended to have larger relative *ΔΔF* values than those that were non-membrane bound and were within a similar range as benchmarked AHs characterized as membrane binders and sensors (Fig. 5B, top; Table S14). The AHs in our list with greater association to the INM had larger relative *ΔΔF* values than those with weak to no association, further supporting our findings that INM binding relies on the detection of packing defects (Fig. 5B, bottom; Table S14).

**Fig. 5.**
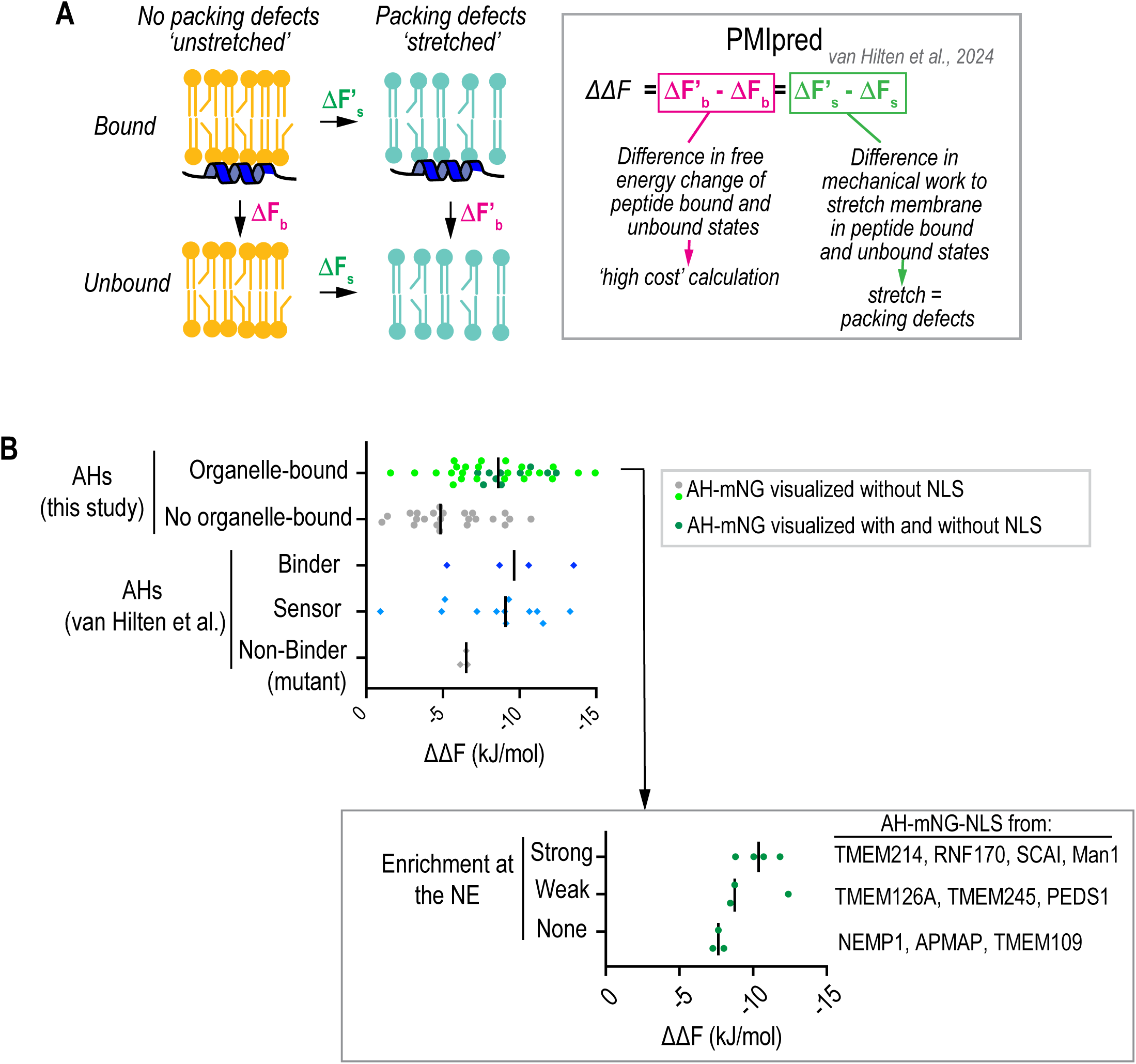
Protein-membrane Interaction predictor (PMIpred) calculations to predict relative sensitivities of AH candidates in NE-associated proteins to packing defects. (A) Schematic representation of how the relative free energy (*ΔΔF*) value is calculated for a given AH as the reduction of work required to ‘stretch’ a leaflet when a peptide is bound. The *ΔΔF* calculation determines the relative change in free energy of a membrane-bound AH with and without ‘stretch,’ which is equivalent to the relative change in free energy of an AH peptide sensing packing defects (van Hilten et al., 2022). (B) Left, *ΔΔF* values of AHs screened in Fig. 1B and benchmark values for known AHs categorized from (van Hilten et al. 2023) predicted by PMIpred (van Hilten et al., 2024). Right, *ΔΔF* values of AHs that were tested in AH-mNG-NLS constructs in Fig. 3. Bar in plots indicates the median value.

Taken together, we conclude that the lipid composition of the INM has adapted to its relatively flat membrane structure by containing lipids that induce packing defects, which can be harnessed by AH sequences in which hydrophobicity through packing defects is the driving force for peripheral membrane association.

We reasoned that because of the minimal curved membranes in the flat, yet loosely packed membrane territory of the INM, AH binding may be exquisitely sensitive to changes in the density of lipid packing defects that result from membrane tension (Pinot et al., 2018; van Hilten et al., 2020). We first used a custom membrane stretch device to impose equibiaxial stretch on living HeLa cells (Kosmalska et al., 2015) to test forces induced by cell-substrate interactions that are known to transmit to cytoskeletal forces imposed directly on the NE. Cell-substrate induced forces can have profound downstream effects on gene expression profiles and cell fate (Carley et al., 2021; Kalukula et al., 2022; Miroshnikova and Wickström, 2022; Nava et al., 2020). These forces mechanically modulate NPCs (Andreu et al., 2022; Elosegui-Artola et al., 2017; Lombardi et al., 2011); however, the effect of mechanical stretch directly on INM tension has not been tested.

We expressed a characterized AH (the ALPS motif) from ArfGAP1 that is highly sensitive to membrane packing defects promoted by membrane tension (Pinot et al., 2018; Shen et al., 2022a). We also chose an AH we identified in TMEM214 protein (Fig. 1B; Fig. S7 and Fig. S11A) because it filled the following criteria: i) localization to the Golgi apparatus when expressed in cytosol (Fig. 1B; Fig. S7) and enrichment of Ser/Thr residues and bulky hydrophobic residues (Fig. S11B) is reminiscent of ALPS motifs (Drin et al., 2007; Magdeleine et al., 2016); ii) predicted to have a *ΔΔF* value (−8.87) that fell in the range of sensor/binder benchmarked AHs (Fig. 5B).

The maximal cell stretch that could be applied without rupturing the plasma membrane while causing the nucleus to deform (see fold increase of cross-sectional nucleus area and pseudo-colored image overlay in Fig. 6A), did not promote INM association of either the ALPS motif of ArfGAP1 or the TMEM214 AH (Fig. 6A). We conclude that INM tension from cell-substrate stretch does not enhance membrane packing defects – these local forces may not propagate INM tension globally, possibly because of membrane flow from the ER to the NE or the dissipation of forces by NPCs and INM proteins interacting with the underlying lamina.

**Fig. 6.**
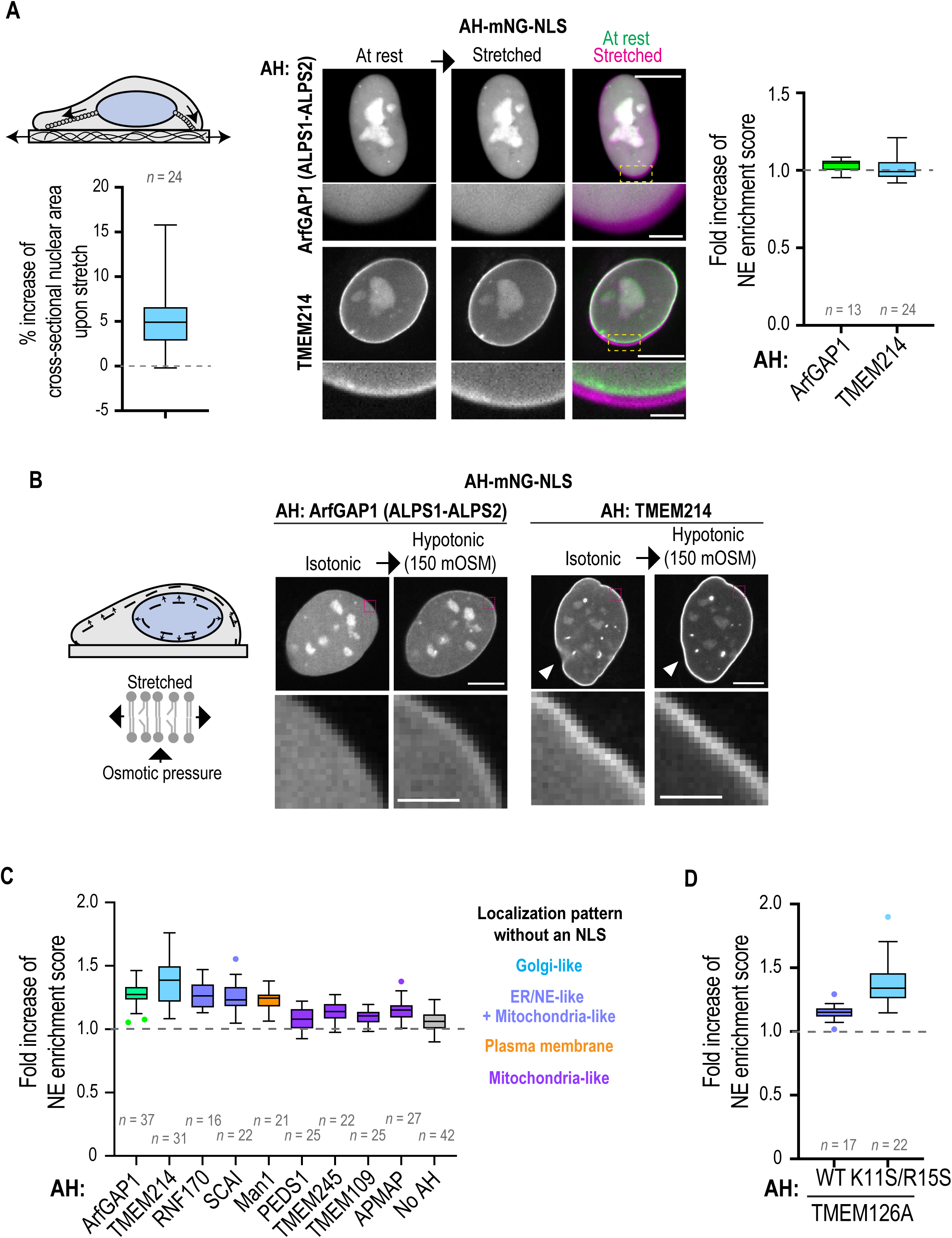
Hypotonic shock and not cell stretch promotes AH binding to the inner nuclear membrane. (A) Schematic representation of nucleo-cytoskeletal linkages directly associated with the nuclear envelope responding to cell-substrate stretch by imposing local forces on the nuclear envelope form the nuclear exterior. Left, Box plot of percentage increase in the cross-sectional nuclear area upon application of cell-substrate stretch. Right, Spinning disk confocal images of single z-section of living HeLa cells expressing indicated AH-mNG-NLS construct before (at rest) and during cell-substrate stretch (stretched). Overlay of pseudo-colored images of nuclei before (green) or during (magenta) cell-substrate stretch shown. Box plot shows fold increase of NE enrichment scores after cell-substrate stretch. Data are pooled from two or three experiments. (B) Schematic representation of uniform internal nuclear forces imposed on the inner nuclear membrane following hypotonic shock increasing tension on the lipid leaflets. Single z-section spinning disk confocal images of living U2OS cells with indicated AH-mNG-NLS before and after hypotonic shock. Arrowhead indicates nuclear rim wrinkle that is no longer present after hypotonic shock. (C and D) Box plot representing calculated fold increase in NE enrichment scores of indicated AH-mNG-NLS before and after hypotonic shock. Data are pooled from two independent experiments. Also see Fig. S12. Scale bars, 10 µm or 2 µm (magnified images).

While local forces may not propagate membrane tension across the NE, we predicted that global INM tension would occur in cells treated with hypotonic shock. Hypotonic shock occurs in various pathophysiological situations such as ischemia and tissue injury and has been shown to promote INM association of cPLA2 (Enyedi et al., 2016; Shen et al., 2022a). Indeed, INM association of the ALPS motif of ArfGAP1 and the AH of TMEM214 was enhanced in U2OS cells treated with mild hypotonic shock (∼150 mOSM) (Fig. 6B and 6C). This is within the range that promotes INM association of cPLA2 (Enyedi et al., 2016) but does not induce organellar vesiculation (King et al., 2020; Niu et al., 2024). AHs from NE proteins had different degrees of INM enrichment in response to mild hypotonic shock, which correlated with their preferred lipid territories when expressed in the cytosol (Fig. 6C; Fig. S12). The K11S/R15S mutant of AH(TMEM126A) displayed a greater response to hypotonic shock as compared to the wild-type AH, supporting the idea that INM-association and membrane tension-sensing by AHs share the detection of packing defects as the underlying mechanism (Fig. 6D; Fig. S12).

Together, these results suggest that membrane tension-sensing is highly prevalent across AHs in NE proteins that are characterized by a paucity of charged residues in the hydrophilic face – for these AHs, the driving force for membrane binding is hydrophobic interactions suggesting that global membrane tension induction by hypotonic shock increases lipid packing defects. Our data also suggest that local forces produced from the cytoskeleton do not propagate across the NE to induce lipid packing defects in the INM. We conclude that different forces should be treated as highly distinct in their consequences on nuclear membrane lipids, which has important implications for how the nucleus responds to specific mechanical inputs.

## Discussion

We have generated a curated and comprehensive list of characterized and uncharacterized transmembrane-containing NE proteins and further analyzed whether they contain predicted AHs. We validated that AHs from NE-associated proteins localize to membrane-bound organelles when expressed in the cytosol and characterized their INM binding propensities. The curated list includes proteins with Ahs that are mutated in diseases associated with the NE (e.g., TMEM214) (Meinke et al., 2020; Reicher et al., 2024; Wilkie et al., 2011), an E3 ligase (e.g., RNF170) (Lu et al., 2011; Song et al., 2020; Wright et al., 2015), and many uncharacterized NE-associated proteins (e.g., TMEM260). By analogy to other membrane-associated AHs in transmembrane proteins, these AHs may regulate the activity or stability (Chua et al., 2017; Halbleib et al., 2017; Kefauver et al., 2020; Lee et al., 2023) or engage in membrane deformation in the context of the full-length protein (Amm et al., 2023; Brady et al., 2015; Breeze et al., 2016; Kayagaki et al., 2021; Kralt et al., 2022). Interestingly, the putative AHs in our study had no overlap with the ALPS-like motifs discovered by (Drin et al., 2007). This could be due to the difference in the AH search algorithm and scope: MemBrain is a deep-learning algorithm with its prediction mechanism being a “black box” that covers a diversity of amino acid sequences that can make up an AH motif (Feng et al., 2022), whereas Drin et al. took a deterministic approach focusing on AHs that belong specifically to the ALPS-motif group (Drin et al., 2007). Thus, our resource complements the existing knowledge of membrane-sensing mechanisms by uncovering novel AHs. Our study of AHs in relation to the NE further suggests that AHs can serve as sensors of membrane tension, which is highly relevant to mechanisms involving mechanosensing and mechanotransduction at the NE.

Inner nuclear membrane-association of the ALPS motif of ArfGAP1, which was shown to be sensitive to membrane tension *in vitro* and *in silico* (Pinot et al., 2018; Shen et al., 2022a), and AHs of INM proteins in response to hypotonic shock suggests that these AHs are sensitive to lipid packing defects resulting from INM tension (Pinot et al., 2018; van Hilten et al., 2020). The fact that tension-sensitive AHs did not bind to the INM upon equibiaxial cellular stretch indicates that the extent of the effect on packing defects depends on the type of mechanical input imposed on the NE. It is established that nuclear deformation upon cellular stretch affects nucleocytoplasmic transport by mechanical modulation of NPCs (Andreu et al., 2022; Elosegui-Artola et al., 2017). Thus, in contrast to hypotonic shock, in which membrane tension is induced by internal forces uniformly imposed on the INM, external cytoskeletal forces on the nucleus induced by cellular stretch only imposes membrane tension locally. These external forces may further be dissipated through tension on NPCs and the underlying nuclear lamina (Carley et al., 2021; Taniguchi et al., 2024) to give rise to a distinct mechanical response.

Our in-depth analysis of AHs in NE-associated proteins suggests that the INM contains packing defects, even under basal conditions. This membrane property may help accommodate nuclear membrane remodeling and repair (Bahmanyar and Schlieker, 2020; Penfield et al., 2020). Additionally, the lipid content of the NE can be altered by both local and global lipid metabolism (Barbosa et al., 2019; Haider et al., 2018; Lee et al., 2023; Romanauska and Köhler, 2018; Romanauska and Köhler, 2021; Samardak et al., 2024; Sołtysik et al., 2021; Tsuji et al., 2019), which has downstream consequences, including local protein degradation (Lee et al., 2023), nuclear growth (Foo et al., 2023; Mauro et al., 2022), and changes in nuclear fragility (Baird et al., 2023; Romanauska and Köhler, 2023).

Our study focuses on lipid packing defect detection by AHs, but our identification of AHs in INM proteins with charged amino acid residues on the hydrophilic face, although correlative, fits with the possibility that low electrostatics may also be present at the INM. However, these AHs did not bind the INM under basal conditions. One possibility is that lipids with negatively charged headgroups are more enriched in specific cell types or are regulated under certain conditions. Indeed, phosphatidic acid (PA) is present at the INM in fungi under specific circumstances and, interestingly, is detected by the AH of the membrane remodeling protein Chmp7 (Garcia et al., 2022; Romanauska and Köhler, 2018; Thaller et al., 2021). Some mammalian cell types can contain PS at the INM; thus, there is more to learn about the role of electrostatics at the INM (Niu et al., 2024; Tsuji et al., 2019). In sum, our work reveals the rules for INM-lipid interactions at the NE by AHs, with broad implications for how the biophysical properties of the INM control NE-dependent functions in both healthy conditions and in diseases associated with the NE.

## Acknowledgments

We thank J.M. Gendron (Yale University, New Haven, CT, USA), Z. Shen and P. Niethammer (Memorial Sloan Kettering Cancer Center, New York, NY, USA), T. Nishimura (Osaka University, Osaka, Japan) and T. Tsuchiya (Tsukuba University, Ibaraki, Japan) for technical advice; D. Gadella (University of Amsterdam, Amsterdam, the Netherlands), M. Davidson (Florida State University, Tallahassee, FL, USA), L.T. Guo and A.M. Pyle (Yale University, New Haven, CT, USA), H. Arai (University of Tokyo, Tokyo, Japan) and Philipp Niethammer for reagents. We thank F. Moore (Yale University, New Haven, CT, USA) for comments on the manuscript. We thank Yale Nucleus Club and BMB Club for helpful discussions. This work was supported by: NIH R01 (GM131004) to S.B.; Swiss National Science Foundation (grant CRSII5_189996), and the European Research Council under the European Union’s Horizon 2020 research and innovation program (grant agreement no. 803952) to S.V.; The Spanish Ministry of Science and Innovation (PID2022-142672NB-I00), the European Research Council (grant 101097753 MechanoSynth), the Generalitat de Catalunya (2017-SGR-1602), the prize “ICREA Academia” for excellence in research to P.R.C. IBEC is a recipient of a Severo Ochoa Award of Excellence from MINCIN.

## Author contributions

S. Lee and S. Bahmanyar conceived the project. S. Lee conceived and performed most of the experiments and data analysis. A.L. Le Roux and P. Roca-Cusachs conceived experiments on cellular stretch and interpreted data. A.L. Le Roux performed cellular stretch experiments. M. Mors performed ΔΔF analysis and was supervised by S. Vanni. S. Lee and S. Bahmanyar wrote the manuscript with input from other authors. S. Bahmanyar supervised the project.

## Lead Contact

Requests for further information, resources, and reagents should be directed to Shirin Bahmanyar.

## Materials Availability

Materials generated in this study are available upon request to the Lead Contact.

## Data and Code Availability

Raw data generated in this study are available upon request to the Lead Contact. Python scripts are found at GitHub as mentioned in Materials and Methods.

## Materials and Methods

### Reagents and resources

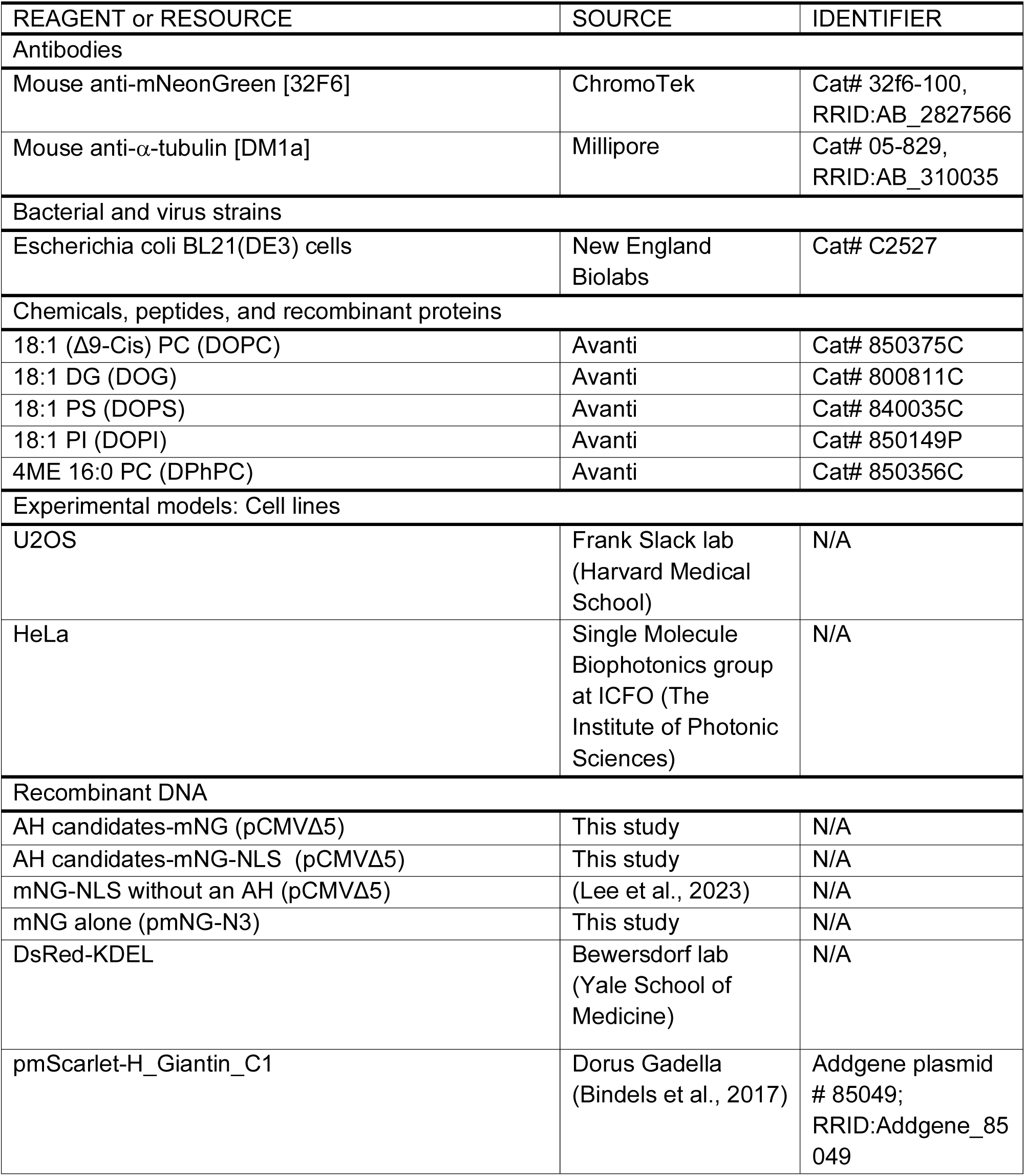
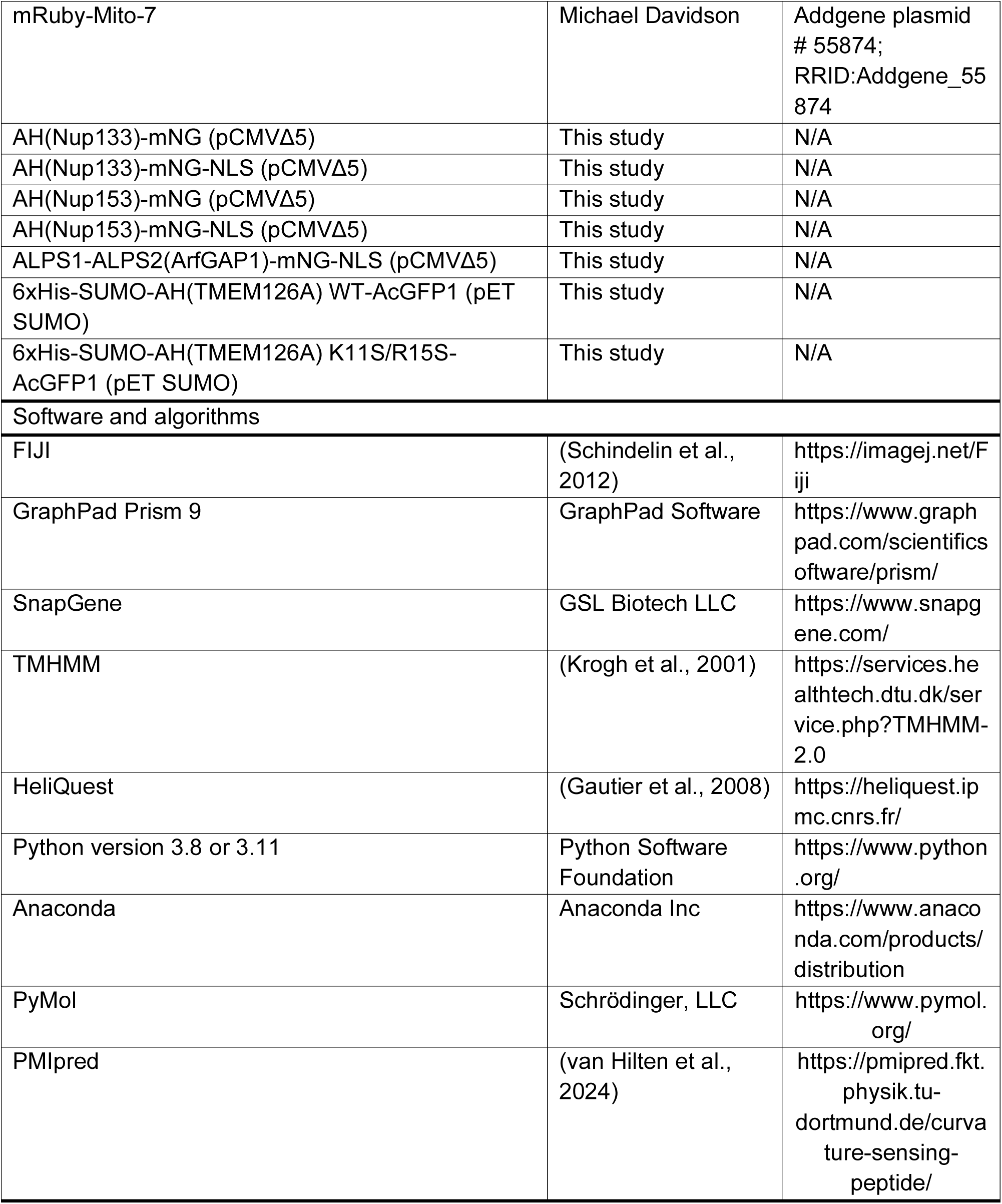

### Computational screening

#### Data mining to generate a curated list of established and putative nuclear envelope-associated proteins

The process described below was performed by custom Python scripts found on Github (https://github.com/shokenlee/Nuclear_proteome). To generate a list of NE-associated proteins, we merged the published dataset from 5 nuclear envelope proteomics papers (Cheng et al., 2019; Korfali et al., 2010; Korfali et al., 2012; Schirmer et al., 2003; Wilkie et al., 2011), using the UniProtID of the human proteins as the unique identifier. First, we first obtained the UniProtID for each protein if it was not available in the original dataset. Listed NCBI accession numbers (Schirmer et al., 2003) or gene names (Korfali et al., 2010; Korfali et al., 2012; Wilkie et al., 2011) were converted to UniProt ID using Retrieve/ID mapping or API of UniProt, respectively. Obsolete or invalid gene records were automatically eliminated. Next, for mouse or rat proteins depending on the study, we retrieved human homologues from UniProt by querying the gene name in humans via API (e.g., the human orthologue of the mouse Ctdnep1 is found by querying “gene name: ctdnep1 & organism: Homo sapiens”). Although this method was not as stringent as an amino acid sequence-based homology search, we leveraged the fact that human, mouse, and rat genes have the same names for homologous proteins in the majority cases. Finally, the five NE proteome datasets were merged to make a final list. In the course of finalizing our screen, (Cheng et al., 2023) published an additional dataset generated from proteomics on the NE so it is not included in our list.

To curate a list of uncharacterized and putative NE-associated proteins, we used data from the Human Protein Atlas (Table S6 in (Thul et al., 2017)) to mine for nuclear proteins based on the following criteria: 1) “IF location score” contains any word including “nucle” (e.g. ‘<u>nucle</u>ar membrane’, ‘<u>nucle</u>oli’); 2) “Reliability” being either “Supported” or “Validated.” For “IF location score”, we included proteins localized anywhere within the nucleus -encompassing not only the nuclear membrane but also nucleoplasm, nuclear bodies, etc. We adopted this broad criterion because 1) The MemBrain list that we used to identify AHs is exclusively composed of putative transmembrane containing proteins, 2) certain nuclear envelope-associated proteins have a soluble pool (e.g., lamin A/C and Lap2α) and/or show nuclear rim and nuceloplasmic localization by immunofluorescence of fixed cells (e.g. LBR in (Lee et al. 2023)), 3) our data mining screening platform was designed to permit false positives while minimizing false negatives.

For secondary analysis, we collected UniProt subcellular location data for the 410 NE proteins through the API. Each protein was considered a NE/ER protein if all of the following three criteria were met: 1) Subcellular location information contained any of the following [’Nucleus outer membrane’, ‘Nucleus membrane’, ‘Nucleus inner membrane’, ‘Nucleus, nuclear pore complex’, ‘Nucleus envelope’, ‘Nucleus lamina’, ‘Endoplasmic reticulum membrane’, ‘Endoplasmic reticulum’, ‘Sarcoplasmic reticulum membrane’, ‘Endoplasmic reticulum-Golgi intermediate compartment membrane’, ‘Endoplasmic reticulum lumen’]; 2) The evidence was manually, but not automatically, curated information. Specifically, the evidence code was any of the following: [’ECO:0000269’ (experimental evidence), ‘ECO:0000305’ (inference from paper), ‘ECO:0000250’ (seq similarity), ‘ECO:0000255’ (seq model), ‘ECO:0000312’ (imported from other database), ‘ECO:0007744’ (a combination of experimental and computational evidence)]; 3) The evidence was not solely reliant on any of the 5 NE proteome papers (PMIDs: [’12958361’, ‘20693407’, ‘20876400’, ‘22990521’, ‘31142202’]) to eliminate duplications. Our rationale to include ER localization as well as NE localization is because the ER and NE share a single membrane with the likelihood that some ER proteins reach the INM. Furthermore, transmembrane containing NE proteins also have a pool in the ER and this subcellular localization is further enhanced by overexpression. Note that although UniProt generally relies on published research, we noticed that some of the NE proteome studies are not reflected in the subcellular location information in UniProt (e.g., TMEM165, TMEM70 (see Table S7)). Consequently, NE proteome studies and UniProt are complementary.

Finally, the NE protein list with UniProt subcellular location data was merged with the Human Protein Atlas list using UniProtID as the unique identifier. Each protein was considered a NE protein if 1) it appeared in 2 or more NE proteome paper, or, 2) it appeared in only one NE proteome paper but was a nuclear protein in Human Protein Atlas or a NE/ER protein in UniProt.

#### Processing and merging the MemBrain list with our curated ‘NE protein list’

The Python scripts for the following are found on GitHub (https://github.com/shokenlee/Find-AH-NE_MemBrain). The MemBrain prediction results for 11,759 proteins from organisms across phylogeny were downloaded from http://www.csbio.sjtu.edu.cn/bioinf/MemBrain/Download.htm (Feng et al., 2022). For each protein, the protein/gene names and organism names were obtained from UniProt through API. Then, the lineage information for each organism was obtained from UniProt through API, and the protein list was screened for those from metazoans. Among them, attempts were made to map non-human proteins to human proteins by searching for the same name of the gene in humans, as conducted for the NE protein listing (see above). Finally, the MemBrain list was merged with the NE protein list using the UniProtID as the unique identifier.

#### Manual assessment of the AH candidates

Secondary analysis of AH candidates was conducted by: 1) manually entering each sequence in HeliQuest to determine the amino acid compositions of hydrophilic and hydrophobic face as well as the mean hydrophobic moment and D factor (Gautier et al., 2008); 2) assessing if amino acid sequence forms a helix by secondary structure prediction using AlphaFold2 (Jumper et al., 2021). An AH candidate was removed if over half of its length overlapped with a juxtaposed transmembrane domain in the context of the full-length protein.; (3) determining if candidate AH sequence is predicted to be a transmembrane helix by inputting into TMHMM (Krogh et al., 2001). Note that a subset of partially membrane-passing AHs were included in the final list because such AHs can be functionally relevant, as exemplified in an ER stress sensor IRE1 where an AH that partially overlaps with the transmembrane helix serves as a membrane stress sensor (Halbleib et al., 2017).

### Plasmid construction

#### General note

Insertion of gene sequences were conducted by using In-Fusion HD Cloning Plus (638909; Takara) or Snap Assembly Master Mix (638948; Takara) unless otherwise noted. Site-directed mutagenesis was performed by whole-plasmid PCR followed by circularization. Successful cloning was confirmed by DNA sequencing all constructs. A bipartite nuclear localization signal (NLS) from Xenopus laevis nucleoplasmin (KRPAATKKAGQAKKKK) (Dingwall et al., 1988; Robbins et al., 1991) was used where indicated.

#### mNG alone construct

The ‘mNG alone’ construct was generated by replacing EGFP in pEGFP-N3 with the mNG sequence (cDNA of mNG encoded in pEGFP-C2 was a gift from Hiroyuki Arai). Note that mNG is expressed under the full-length pCMV promoter.

#### Cloning of candidate AHs tagged with mNG

A pool of single strand DNA oligos encoding candidate AHs were synthesized by Twist Biosciences. The sequences included <u>AAACGGGCCCTCTAGAGCC</u>ACC*ATG*GGA (GGA)n XXX (GGT)n GGAT<u>CGAATTCTGCAGTCGACGGTAC</u>, where XXX indicates each AH sequence, underlined sequences indicate the annealing sites for primers, *ATG* is the start codon, (GGA)n and (GGT)n are glycine linkers with n greater than or equal to zero and less than or equal to 7. The length of glycine linkers was chosen to keep the length variance of DNA oligos in the pool within +/- 10% of the median length for better efficiency of oligo pool synthesis, as instructed by Twist Biosciences. (2) PCR was performed with the single stranded oligo pool used as a template by forward and reverse primers (complementary to underlined sequences indicated above) with KAPA HiFi HotStart ReadyMix PCR Kit (50-196-5217; Roche Diagnostics) and purified with NucleoSpin Gel and PCR Clean-Up (740609; Takara). (3) Purified amplicons were inserted into a linearized plasmid with In-Fusion HD Cloning Plus (638909; Takara). The backbone plasmid (pCMVΔ5-Sun2-mNG; (Lee et al. 2023)), where the mNG sequence was taken from the ‘mNG alone construct’ in pEGFP-N3, was linearized by inverse whole-plasmid PCR, removing the coding sequence of Sun2. The plasmid has a clipped CMV promoter (CMVΔ5) (Morita et al., 2012) to minimize the expression level without compromising transfection efficacy. The resulting plasmid has the glycine linker introduced from the pooled oligos and an amino acid sequence “NSAVDGTAGPGSAT” linker after the AH sequence originating from the multiple cloning site of pEGFP-N3 followed by the mNG sequence. Approximately 120 colonies were picked and mini-prepped to obtain the 57 AH-mNG constructs.

#### Cloning of AH sequences of Nup133, Nup153 and the ALPS motifs of ArfGAP1 tagged with mNG

The ALPS-like motif (residues 245-263) of human Nup133 (UniProt ID Q8WUM0; (Drin et al., 2007)) was synthesized and PCR-amplified. The AH (residues 35-62) of human Nup153 (UniProt ID P49790) was amplified from pEGFP3-Nup153 (Daigle et al., 2001). The AH (ALPS1-ALPS2) of ArfGAP1 was amplified from pGEX-6P-1-hs-ALPS1-ALPS2-mCherry (a gift from Philipp Niethammer (Addgene plasmid # 187114; RRID: Addgene_187114); (Shen et al., 2022b)). The amplicons were inserted into the same backbone plasmid as the AH candidates with and without the sequence encoding the nucleoplasmin NLS.

#### His-SUMO-AH-AcGFP1

AcGFP1 was amplified from GFP-Sec61b (a gift from Gia Voeltz) and inserted into the pET-SUMO vector (ThermoFisher Cat# K30001). The AH DNA sequence was amplified from the corresponding AH-mNG construct and inserted in between SUMO and AcGFP1 with “RQAMGG” and “GGSNSAVDGTAGPGSAT” linkers being before and after the AH, respectively. AcGFP1 has been shown to be monomeric *in vitro* (Gurskaya et al., 2003) .

### Tissue culture

U2OS cells were grown at 37°C in 5% CO_2_ in DMEM low glucose (11885; Gibco) supplemented with 10% heat inactivated FBS (F4135) and 1% antibiotic-antimycotic (15240112; Gibco). HeLa cells were grown in DMEM (Thermo Fisher Scientific; 11960-044) supplemented with 10% FBS (Thermo Fisher Scientific; 10270-106), 1 mM sodium pyruvate (Thermo Fisher Scientific; 11360039) and 1% penicillin-streptomycin (Thermo Fischer Scientific; 10378-016). Cells were used for experiments before reaching passage 25. Cells were continuously profiled for contamination by assessment of extranuclear SiR-DNA staining.

### Transfection

U2OS cells were seeded to reach 50–80% density on the day of transfection. DNA transfections were performed with Lipofectamine2000 (11668; Thermo Fisher Scientific) in Opti-MEM (31985; Gibco) with DNA concentrations ranging from 50 to 300 ng DNA per cm^2^ of growth surface. HeLa cells were transfected with 6 µg of plasmid using the Neon transfection device (Thermo Fisher) according to the manufacturer’s instructions and parameter listed for HELA cells (2 pulses of 35 s at 1005 V). In both cases, cells were imaged 24 h after DNA transfection.

### Live-cell imaging

For live imaging, cells were plated in µ-Slide 8-well Glass Bottom chamber (80827; ibidi). Samples were imaged in a CO2-, temperature-, and humidity-controlled Tokai Hit Stage Top Incubator. The imaging media used was DMEM supplemented with 10% FBS and 1% antibiotic-antimycotic (15240112; Gibco). For SiR-DNA staining, cells were incubated with 250 nM SiR-DNA (Lukinavičius et al., 2015; Sen et al., 2018) in the growth media for 1 hr prior to imaging.

### Microscopy

Live-cell imaging was also performed on an inverted Nikon Ti Eclipse microscope equipped with a Yokogawa CSU-W1 confocal scanner unit with solid state 100 mW 405, 488, 514, 594, 561, 594, and 640 nm lasers, using a 60× 1.4 NA plan Apo oil immersion objective lens and/or 20× plan Fluor 0.75 NA multi-immersion objective lens, and a prime BSI sCMOS camera. GUV imaging was performed on an inverted Nikon Ti microscope equipped with a Yokogawa CSU-X1 confocal scanner unit with solid state 100mW 488-nm and 50-mW 561-nm lasers, using a 60× 1.4 NA plan Apo oil immersion objective lens, and a Hamamatsu ORCA R-2 Digital CCD Camera.

### Hypotonic treatment

Cells in isotonic condition (i.e., cultured in regular media) were imaged prior to exchanging the culture medium to a mixture of 150 µL of cell culture media and 150 µL of MilliQ water (resulting in approximately 150 mOSM). The same cells were imaged after changing the medium, within ∼3 minutes.

### Cell stretch experiments

#### Preparation of stretchable PDMS membranes

Stretchable polydimethylsiloxane (PDMS) (Sylgard Silicone Elastomer Kit, Dow Corning) membranes were prepared as previously described (Casares et al., 2015; Kosmalska et al., 2015). Briefly, a mix of 10:1 base to crosslinker ratio was degassed and spin-coated for 1 min at 500 rpm and cured at 65 °C overnight on methacrylate plates. Once polymerized, membranes were peeled off and assembled onto a metal ring that can subsequently be assembled in the stretch device.

#### Mechanical stimulation of cells and microscopy for live imaging of stretched cells

Cell mechanical stimulation was done as previously described (Kosmalska et al., 2015) using a custom-built stretching device that produced uniform biaxial deformations of stretchable PDMS membranes and of overlying cells. Briefly, a 150 μL droplet of a 10 μg/mL fibronectin solution (Sigma, F1141) was deposited at the center of the PDMS membrane mounted in the ring. After overnight incubation at 4 °C, the fibronectin solution was rinsed with PBS (Gibco, 14200-067), cells were seeded on the fibronectin-coated membranes and allowed to attach for 30–90 min in presence of Spy650-DNA (1:1000 dilution, Spirochrome, SC501). Before the experiment, the media was exchanged to CO_2_-independent media supplemented with rutin (Bogdanov et al., 2012). Then ring-containing membranes were mounted in the stretch system previously described (Casares et al., 2015; Kosmalska et al., 2015). The stretch system was mounted inside of an inverted microscope (Nikon Eclipse Ti) equipped with an incubation chamber, and the temperature was kept et 37°C during the experiment. Images of cells were acquired with a 60x objective (NIR Apo 60X/WD 2.8, Nikon) on the inverted microscope, with a spinning disk confocal unit (CSU-W1, Yokogawa), a Zyla sCMOS camera (Andor) and using the Micromanager software. For each cell, a z-stack of nuclei was acquired at rest (0.2 μm step) was recorded using the 488 nm excitation wavelength for mNeon-green and 640 nm excitation wavelength for SpyDNA. Several cells (5 - 6) were imaged at rest (before stretching) each stretch experiment and their positions were recorded. After the PDMS substrate was stretched (12 - 15% substrate strain), substrate movement due to stretch resulted in cell movement refocusing followed by imaging with same setting as pre-stretch.

### Image analysis

The “NE enrichment score” was determined based on spatial quantification of nuclear and nuclear rim fluorescence intensities performed on high resolution single plane images. A modified custom ImageJ Macro (Lee et al., 2023) was used to (see also Fig. S13A for the workflow): (a) Auto-segment the SiR-DNA channel by the Li method (Li and Lee, 1993) to define the entire nuclear region as the ROI. (b) Apply the ROI to the mNG channel and measure the integrated fluorescence intensity of the nucleus (“Total nucleus”). (c) Erode the ROI three times and use eroded ROI to mask the mNG channel and measure the “intra-nuclear” integrated fluorescence intensity. (d) Subtract these values to quantify the integrated fluorescence intensity of the “nuclear rim.” (e) Quantify the mean fluorescence intensity values by dividing the integrated fluorescence values of the “Total nucleus” and “Nuclear rim” by their corresponding areas. Area of nuclear rim was determined using the same procedure as for determining the fluorescence intensity. (f) Subtract background mean intensity from each value. (g) Quantify the ratio of the “Nuclear rim” to “Intra-nuclear” mean fluorescence intensity values to obtain the “NE enrichment score.” Pool measurements from multiple cells were plotted using a box plot to display the distribution of values and median.

Quantification of *in vitro* purified AH-AcGFP proteins localization to GUV rims made up of different lipid compositions were based on fluorescence intensity measurements performed on high resolution single plane images using a custom ImageJ macro. GUVs in the red channel were auto-segmented by the Li method to obtain ROI #1 (see Fig. S13B for illustration of ROIs). Then, the segmented image was eroded two times and used to obtain ROI #2. The mean fluorescence of the GFP channel in ROI#2 was subtracted from ROI #1 to obtain the fluorescence intensity value of the “GUV rim”. ROI #2 was then eroded three more times pixels to obtain ROI #3, which was used to obtain a background value. ROI #3 was then dilated 10 and 15 times to define ROIs #4 and #5, respectively. Subtraction of the GFP mean fluorescence intensity in ROI #4 from that in ROI #5 was performed to obtain the GFP mean fluorescence intensity “Outside of GUV” corresponding to unbound GFP protein. The mean fluorescence intensity value from the background was subtracted from “GUV rim” and “Outside GUV” mean fluorescence intensity values. The ratio between “GUV rim” to “Outside of GUV” for each GUV was quantified and pool measurements from multiple GUVs were plotted using a box plot to display the distribution of values and median.

### D factor

D factor was given by: D = 0.944 * µH + 0.33 * z, where µH is the magnitude of hydrophobic moment and z is the net charge of a given AH, accordingly to HeliQuest (https://heliquest.ipmc.cnrs.fr/HelpProcedure.htm “\l” heading3) (Drin et al., 2007; Kardani and Bolhassani, 2021; Su et al., 2022).

### Mapping the hydrophobic and hydrophilic faces of predicted amphipathic helices

The analysis was performed with custom Python scripts (https://github.com/shokenlee/AH_AAcomposition) inspired by HeliQuest (Drin et al., 2007). (1) The amino acid residues were placed in a helical wheel projection with the assumption that all of the AHs are α-helices with 100 degree rotation per amino acid residue (3.6 amino acid residues per turn). (2) To determine the hydrophobic and hydrophilic faces of the helix, the direction (θ) and magnitude (µH) of the mean hydrophobic moment vector were calculated using the standard hydrophobicity scale (Fauchere and Pliska, 1983). Based on the direction (θ) of the mean hydrophobic moment vector, a perpendicular line was drawn across the wheel projection.

### Immunoblot

U2OS cells plated on a 24-well plate were lysed with 50 µL of ice-cold RIPA buffer (25 mM Tris pH 7.4,1% NP-40, 0.5% sodium deoxycholate, 0.1% SDS, 150 mM NaCl,and 1 tablet/50 ml cOmplete Mini protease inhibitor cocktail [11836153001; Roche]), incubated on ice for 15 min, and then centrifuged at >20,000 × g (15,000 rpm) for 15 min at 4°C. Eighteen microliters of whole cell lysate was loaded per lane in a 12% polyacrylamide gels. Proteins were wet-transferred to 0.22 μm nitrocellulose membranes (1620112; Bio-Rad). Nitrocellulose membranes were blocked in 1% BSA in PBS for 30 mins, incubated with primary antibodies diluted in 1% BSA for 1.5 h at room temperature with rocking. Nitrocellulose membranes were incubated with goat anti-mouse IgG secondary antibodies (31430; Thermo Fisher Scientific) in 5% milk in TBST for 45 min at room temperature with rocking. All wash steps were with three times for 5 min in TBST. Clarity ECL reagent (1705060S; Bio-Rad) or ECL select (RPN2235; Cytiva) (in case of low expression level of AH-mNG proteins) was used to visualize chemiluminescence with a Bio-Rad ChemiDoc Imaging System. Antibody concentration was the following: anti-mNG 1:3,000; α-tubulin 1:5,000; secondary antibodies 1:10,000.

### Recombinant protein purification

6xHis-SUMO-AH-AcGFP1 proteins were expressed in Escherichia coli BL21(DE3) cells. Cells were grown at 37 °C to an OD 600nm of 0.4 - 0.7 and then cooled at 20 °C. Protein expression was induced with 0.1 mM isopropyl βD-1-thiogalactopyranoside (IPTG) at 20 °C for 16 h, and cells were harvested by centrifugation and stored at −80 °C. Frozen cells were resuspended in lysis buffer (50 mM Na_2_HPO_4_, 500 mM NaCl, 10 mM imidazole, pH 8.0) and lysed by sonication. After adding betamercaptoethanol to a final concentration of 10 mM, samples were centrifuged at 21,000 g at 4 °C for 20 min. The supernatant was incubated with pre-equilibrated Ni-NTA agarose (Qiagen 30210) at 4 °C for 1 h. The resin was centrifuged at 2,000 rpm at 4 °C for 2 min, washed 3 times with wash buffer (50 mM Na_2_HPO_4_, 300 mM NaCl, 20 mM imidazole, pH 8.0). The His tagged protein was eluted with elution buffer (50 mM Na_2_HPO_4_, 300 mM NaCl, 250 mM imidazole, pH 8.0), aliquoted and analyzed by SDS-PAGE and Coomassie blue stain. Fractions containing the protein were collected and combined. The 6xHis-SUMO tag was cleaved by treatment with recombinant 6xHis-Ulp1 (a gift from Anna Marie Pyle, Yale University; (Guo et al., 2020)) at 4°C for 1 h following incubation with Ni-NTA agarose to remove the 6xHis-SUMO tag and 6xHis-Ulp. AH-AcGFP1 protein was buffer-exchanged with 1xPBS by using Amicon centrifugal filter MWCO 10 kDa (Millipore Sigma Cat# UFC5010), then flash-frozen and stored at −80 °C until used for GUV assays.

### Preparation and imaging of giant unilamellar vesicles (GUVs)

Giant Unilamellar Vesicles (GUVs) were prepared by polyvinyl alcohol (PVA)-assisted swelling (Romanauska and Köhler, 2023; Weinberger et al., 2013). PVA (weight-average molecular weight of 146,000 to 186,000; Cat# 363065; Sigma-Aldrich) was solubilized in water to 5% (w/w) stirred and heated at 90 °C for 1 hr and stored at room temperature until used. Fourty microliters of PVA solution was spread on a glass coverslip (1.2 cm diameter), which was pre-washed with 70% ethanol and distilled water, and dried at 50 °C for 30 min. PVA coated coverslips were stored at room temperature. Lipid mixture in chloroform was prepared in a glass tube (Fisher; #14-961-26) containing 0.25 mol/mol% of Rhodamine-PE to label GUV. Two microliters of lipid mixture (1 mg/ml) dissolved in chloroform was spread on top of the PVA layer and chloroform was allowed to evaporate for 1 h. Two-hundred microliters of swelling solution (280 mM sucrose) was added on each coverslip placed in 24-well cell culture dish and incubated for 1 h at room temperature with gentle rocking to induce vesicle formation. The solution containing vesicles was collected and pelleted by centrifugation at 500 g for 5 min. Vesicles in pellet were suspended in 80 µL of 1xPBS and kept at room temperature for immediate use. Vesicles were mixed with purified proteins (final protein concentration 500 nM) and immediately imaged in an ibidi 8 well chamber on a spinning disk confocal microscope, as described above. Wells were blocked for 1 h with 2.5 mg ml−1 bovine serum albumin in PBS and were washed with 1xPBS before loading the samples.

### In silico prediction of *ΔΔF* values

The affinity of each AH sequence for membranes with increased lipid packing defects, quantified by *ΔΔF* values (van Hilten et al., 2022), was predicted using PMIpred (van Hilten et al., 2024) (https://pmipred.fkt.physik.tu-dortmund.de/curvature-sensing-peptide/) with a neutral membrane as the target. The 19 peptides that serve as comparison were obtained from the benchmark set of van Hilten et al. (2023), where they were experimentally classified as non-binders, binders, or sensors. For this study, their *ΔΔF* values were also calculated with PMIpred.

### Analysis of the ALPS-like motif list

The list of proteins with putative ALPS-like motifs was obtained from the Supplementary Table S1 in (Drin et al., 2007). UniProt IDs in the list were validated in UniProt using API, which resulted in removal of 3 obsolete IDs. The remaining, valid proteins (193 in total) were crossed with our lists of NE proteins and predicted AHs.

### Statistical analysis

GraphPad Prism 8 was used for all statistical analysis otherwise specified in methods. Statistical tests used, sample sizes, definitions of n and N, and p values are reported in Figures and/or Figure legends.

---

**Fig. S1.**
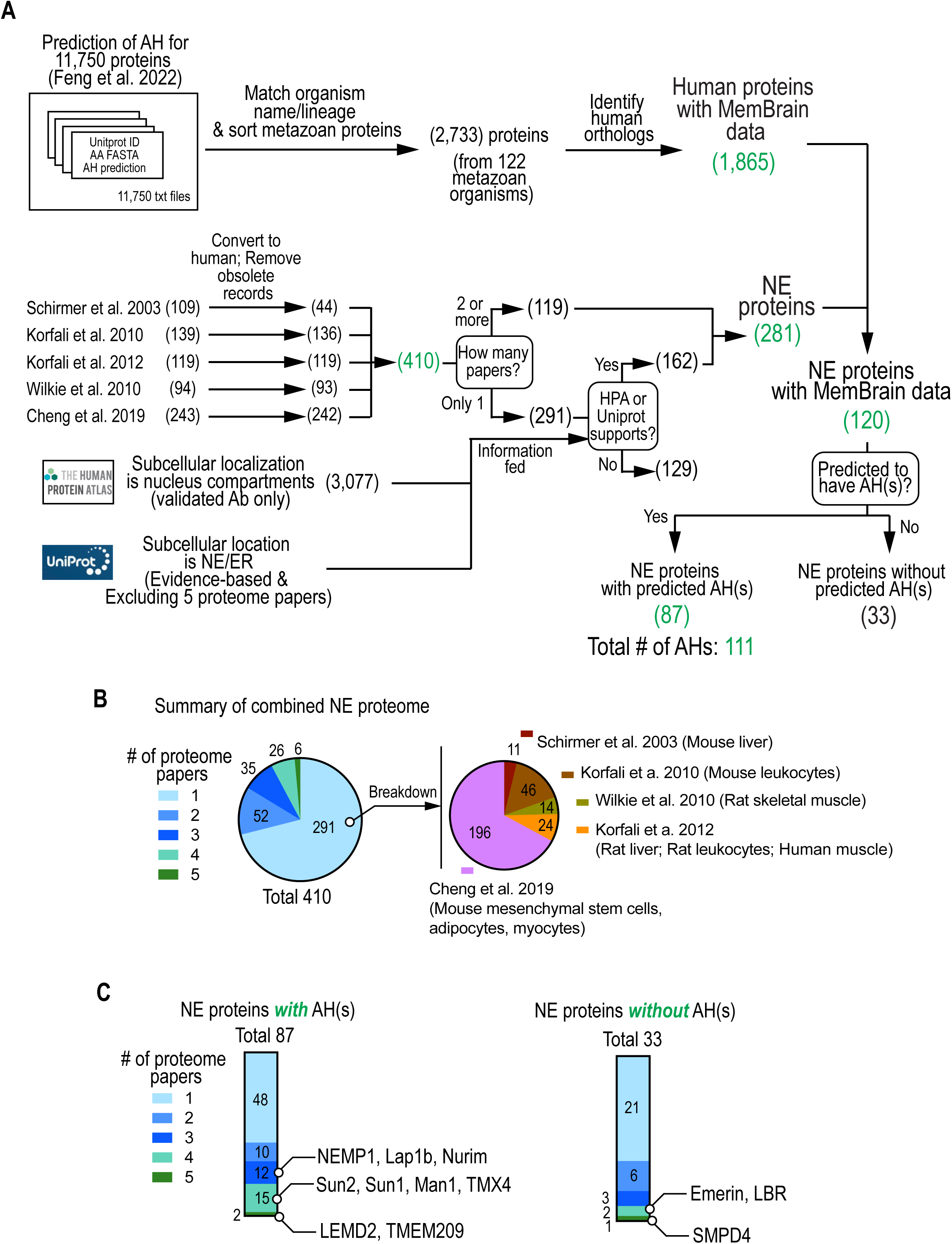
Computational screen of AHs in nuclear envelope proteins. (A) Detailed flow chart of data mining for putative NE proteins and identification of predicted AHs. Numbers in parenthesis indicate the number of proteins. (B) Breakdown of the 410 NE proteins by the number of the NE proteome papers. NE proteins found in only a single NE proteome paper were further categorized by the specific NE proteome paper. (C) Breakdown of NE proteins with or without a predicted AH by the number of NE proteome papers.

**Fig. S2.**
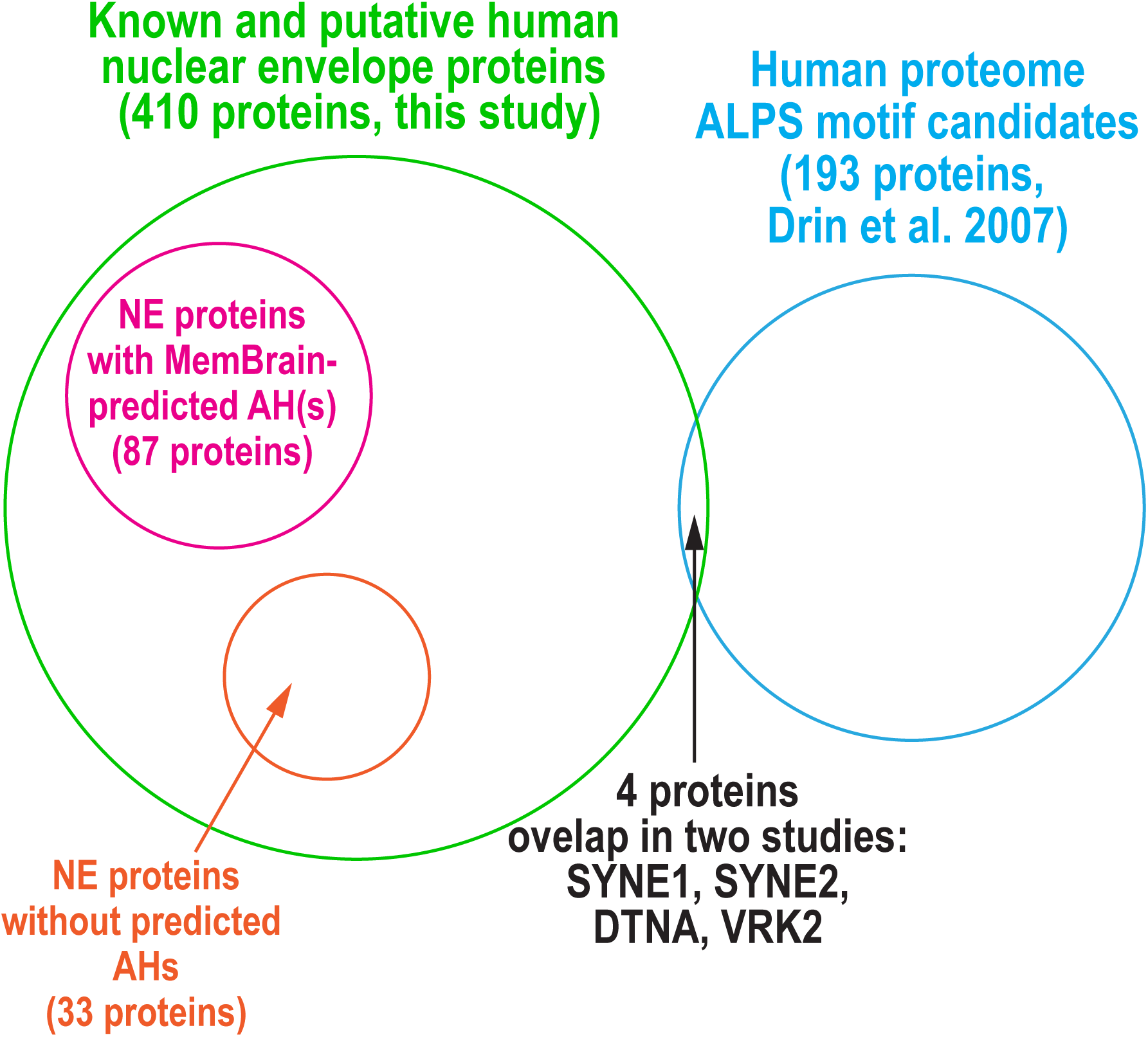
Overlap of AHs in this study and ALPS-like motifs in Drin et al. 2007. Venn diagram representing the NE proteins, those with or without a MemBrain-predicted AH, and proteins with a putative ALPS-like motifs (Drin et al. 2007).

**Fig. S3.**
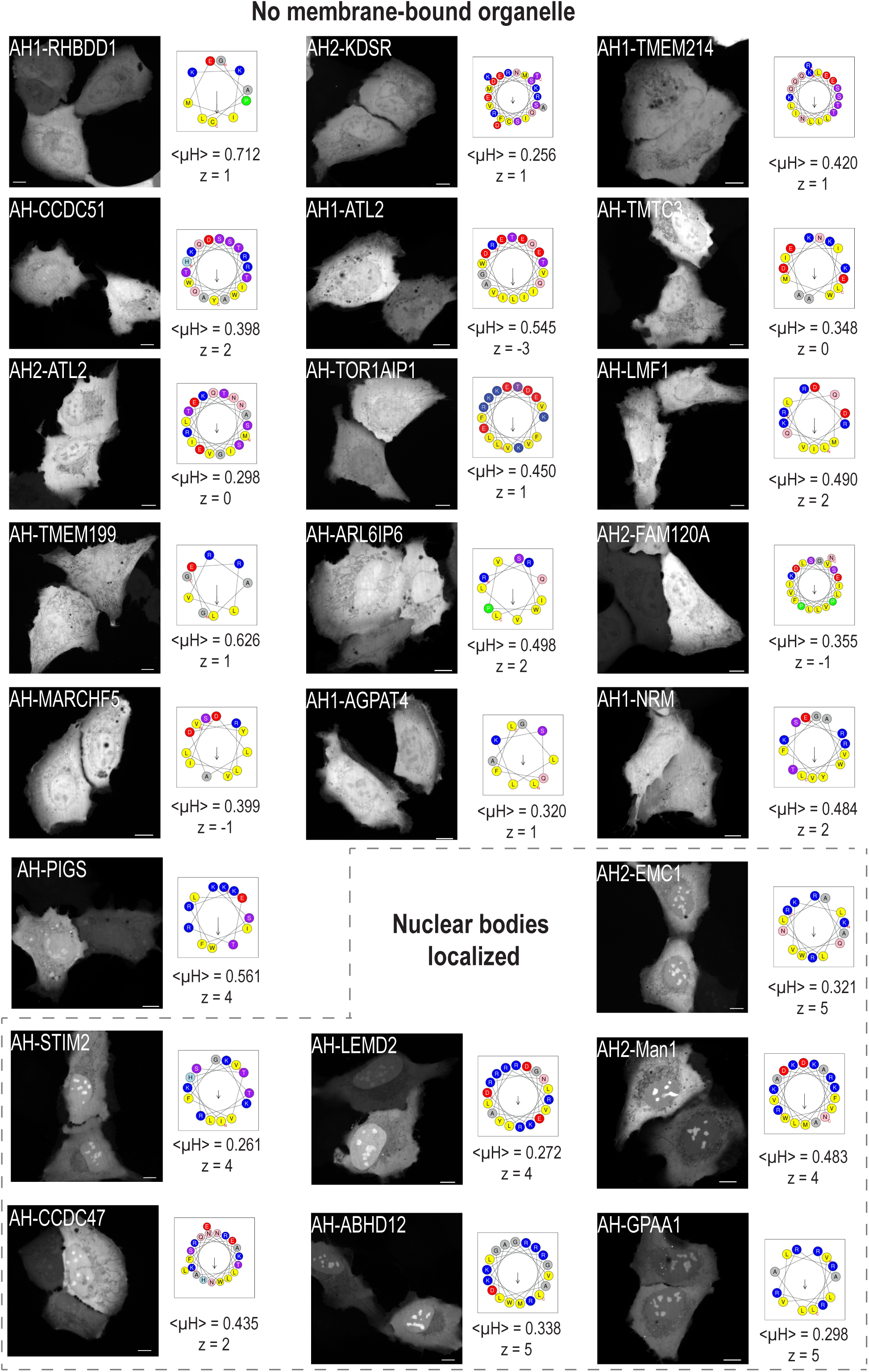
Subcellular localization patterns of AH candidates tagged with mNG that did not localize to membrane-bound organelles. Representative spinning disk confocal images of a single section in living U2OS cells expressing the indicated AH-mNG protein are shown. Schematic representations of corresponding helical wheel projections of the predicted AHs with mean hydrophobic moment <µH> and net charge z is shown. AH-mNG proteins with a nuclear body-like pattern are highlighted at the bottom right. Scale bars, 10 µm.

**Fig. S4.**
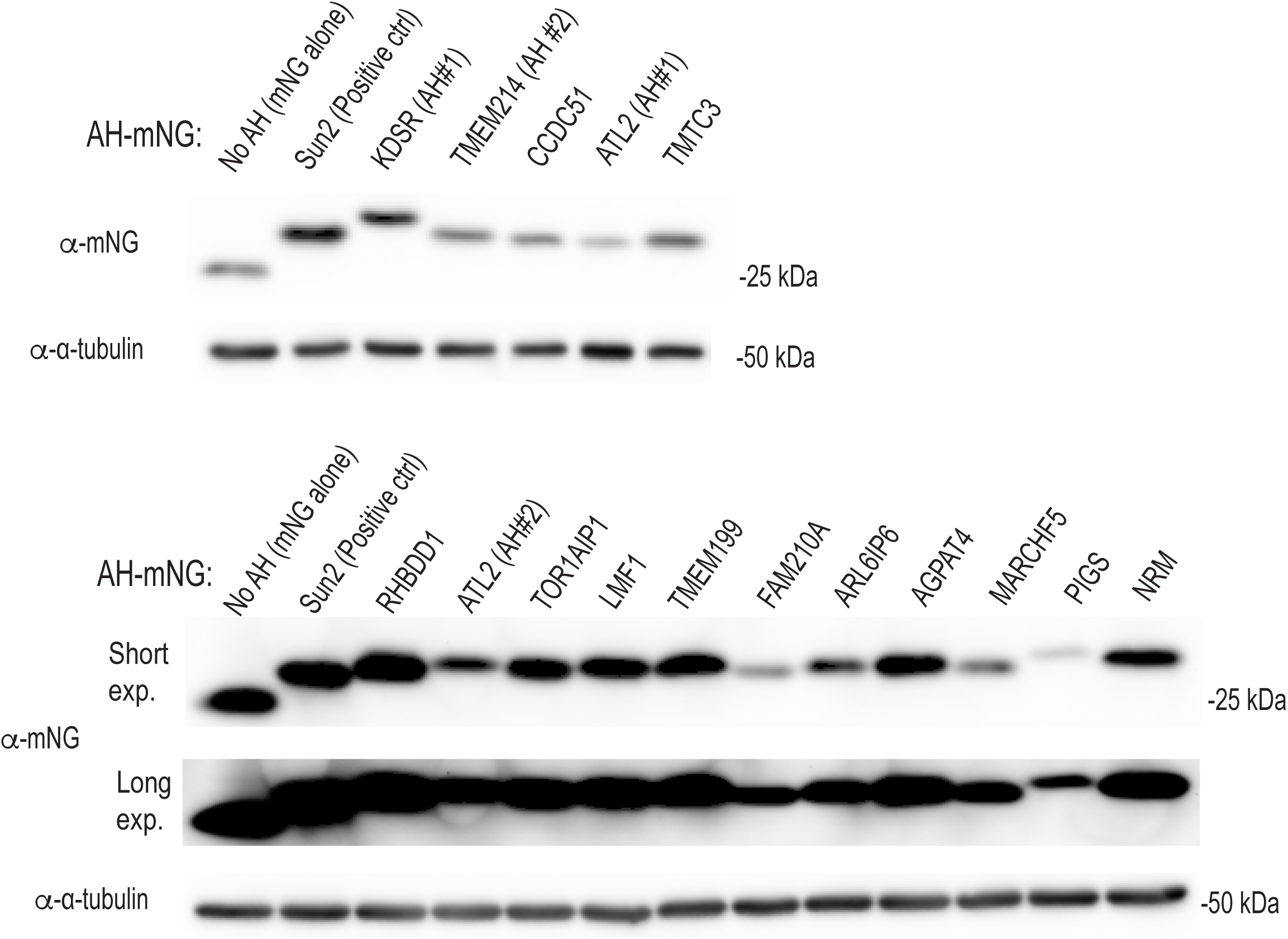
AH-mNG without localization to membrane-bound organelles or nuclear bodies is expressed without unexpected cleavage of the AH. Immunoblot of mNG alone or the indicated AH-mNG proteins transiently expressed in U2OS cells. Among the AH-mNG proteins without membrane-bound organelle localization (Fig. S2), we tested AH-mNG proteins that did not localize to nuclear bodies, because the nuclear body localization indicated the presence of AH in the expressed protein.

**Fig. S5.**
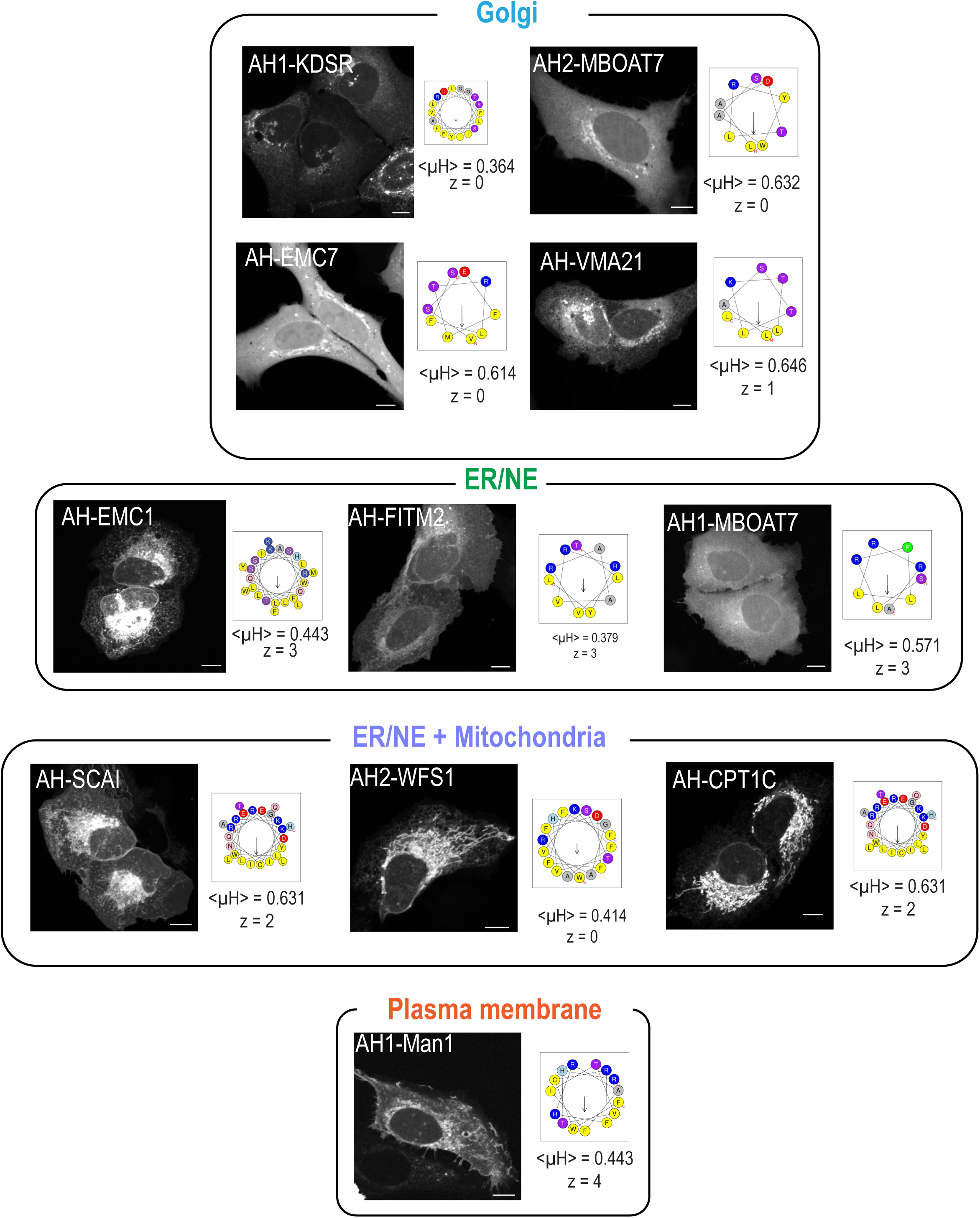
Subcellular localization patterns of AH candidates tagged with mNG that displayed non-mitochondrial patterns. Representative spinning disk confocal images of a single section in living U2OS cells expressing the indicated AH-mNG protein are shown. Schematic representations of corresponding helical wheel projections of the predicted AHs with mean hydrophobic moment <µH> and net charge z is shown. Scale bars, 10 µm.

**Fig. S6.**
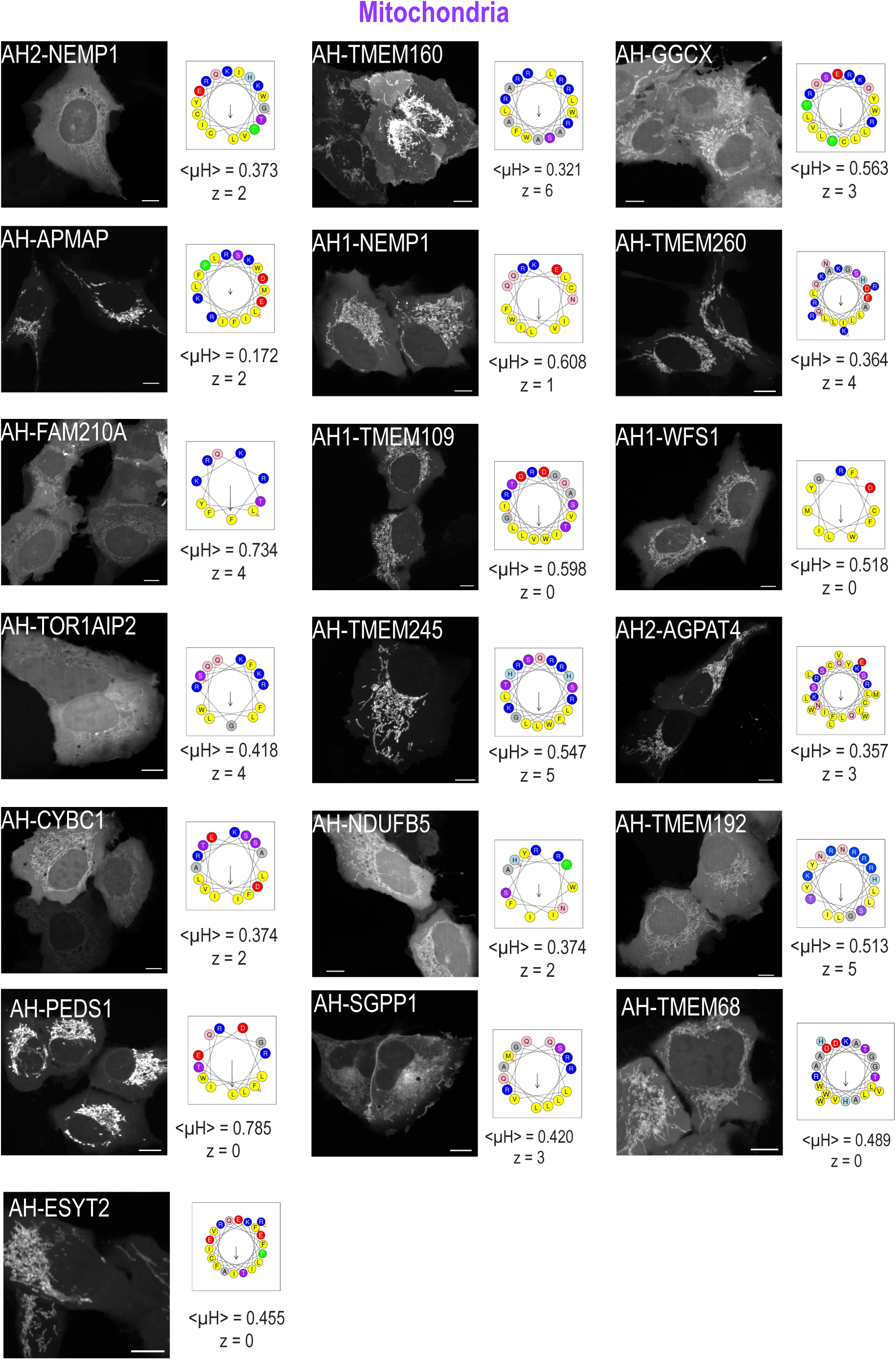
Subcellular localization patterns of AH candidates tagged with mNG that displayed mitochondrial patterns. Representative spinning disk confocal images of a single section in living U2OS cells expressing the indicated AH-mNG protein are shown. Schematic representations of corresponding helical wheel projections of the predicted AHs with mean hydrophobic moment <µH> and net charge z is shown. Scale bars, 10 µm.

**Fig. S7.**
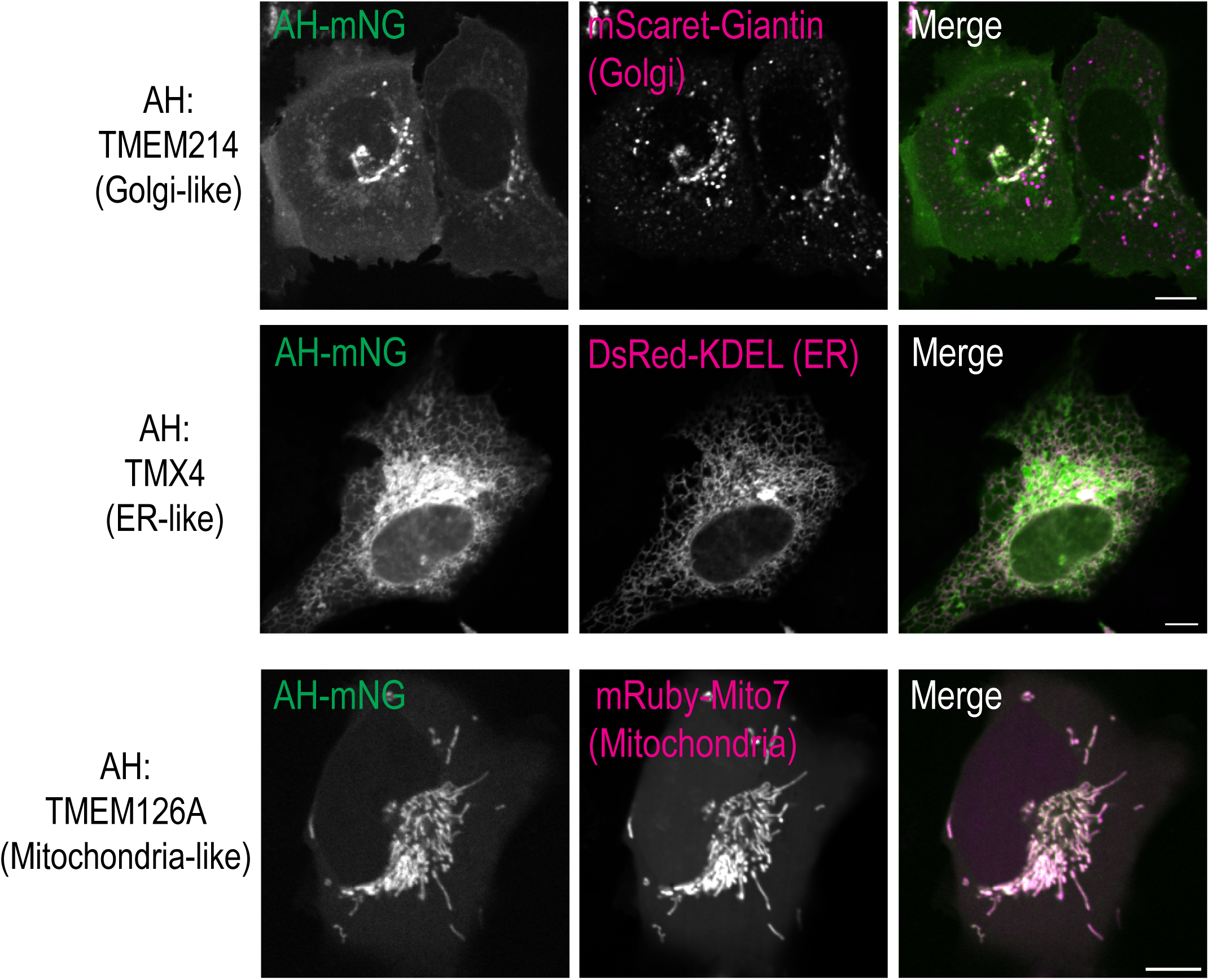
Assessment of the organelle type to which that AH-mNG constructs localized. Representative spinning disk confocal images of a single section in living U2OS cells co-expressing the indicated AH-mNG and the indicated organelle marker are shown. Scale bars, 10 µm.

**Fig. S8.**
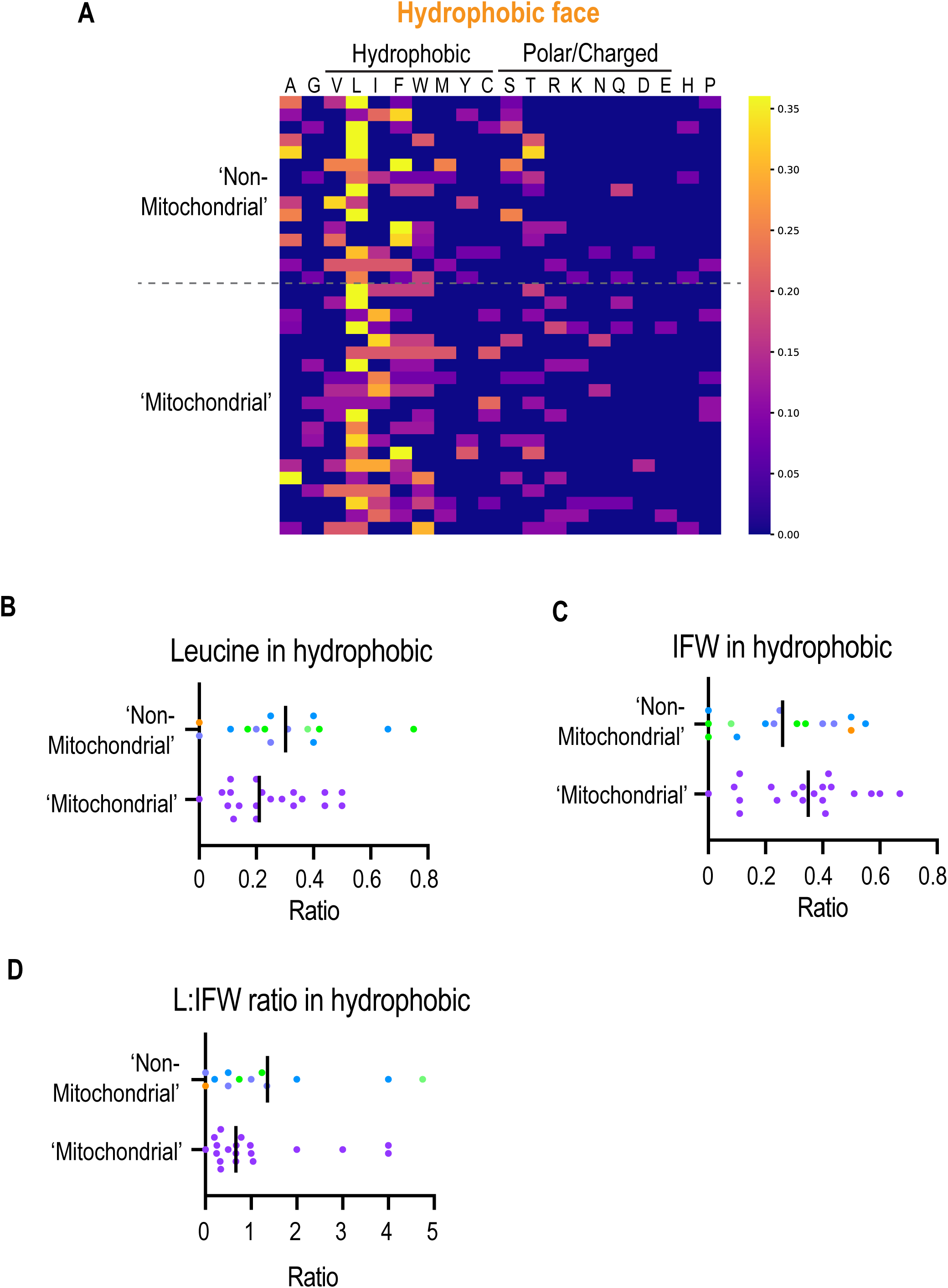
Sequence codes in hydrophobic face that discriminate AHs with non-mitochondrial and mitochondria-like patterns. (A) Heat map representation of the fraction of each amino acid (columns) present within the hydrophobic face of each amphipathic helix (row) categorized by indicated localization patterns. (B-D) Scatter plots of the indicated metrices of AH candidates categorized and color-coded by their localization patterns as determined from experiments in Fig. 1B.

**Fig. S9.**
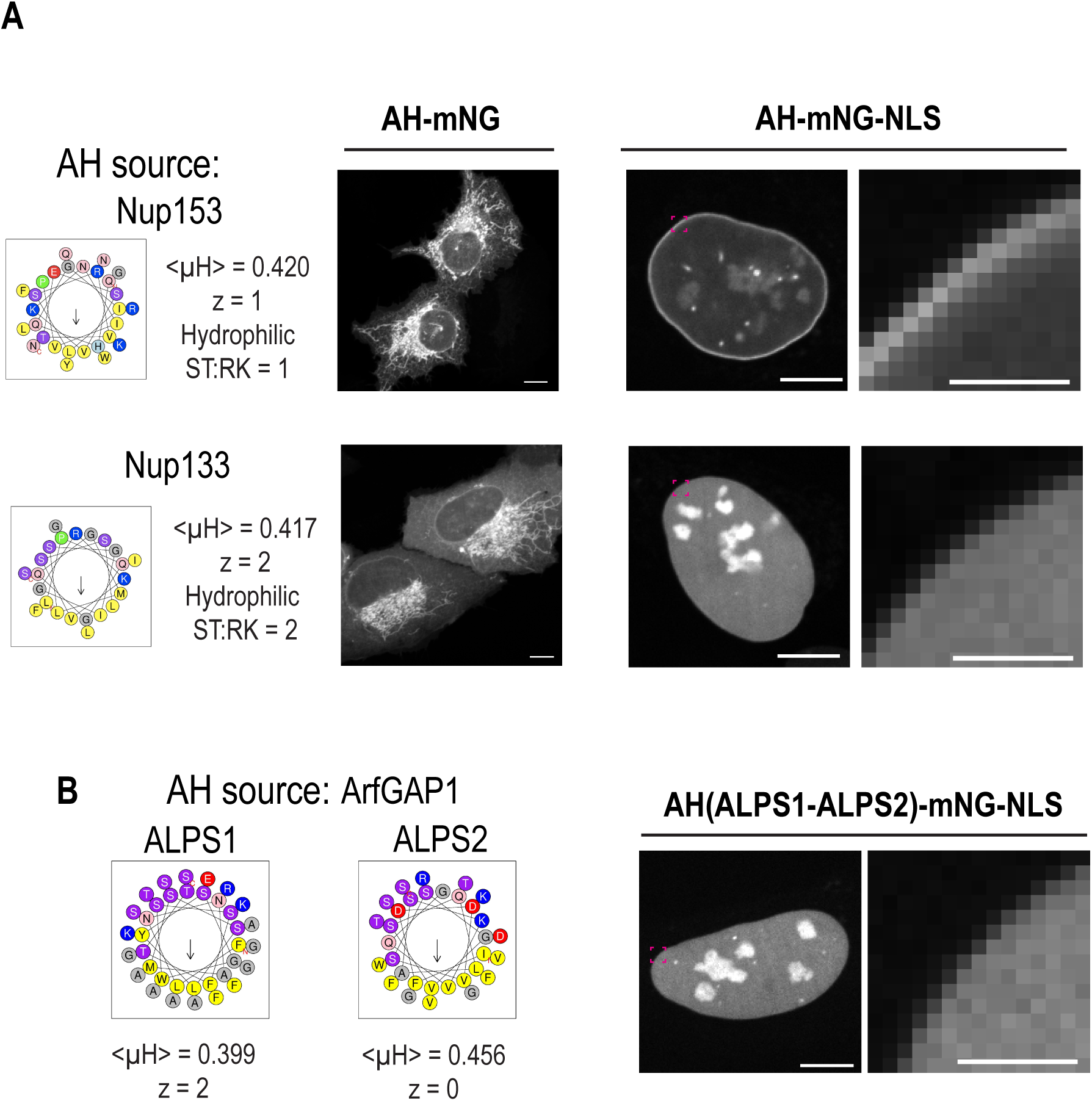
Subcellular localization patterns and INM association of AHs from human Nup153, Nup133 and ArfGAP1. Representative spinning disk confocal images of a single section in living U2OS cells expressing the indicated AH-mNG (A) and AH-mNG-NLS protein with magnified zoom of the nuclear rim area (A and B) are shown. Schematic representations of corresponding helical wheel projections of the AHs with mean hydrophobic moment <µH> and net charge z is shown. Scale bars, 10 µm or 2 µm (zoom)

**Fig. S10.**
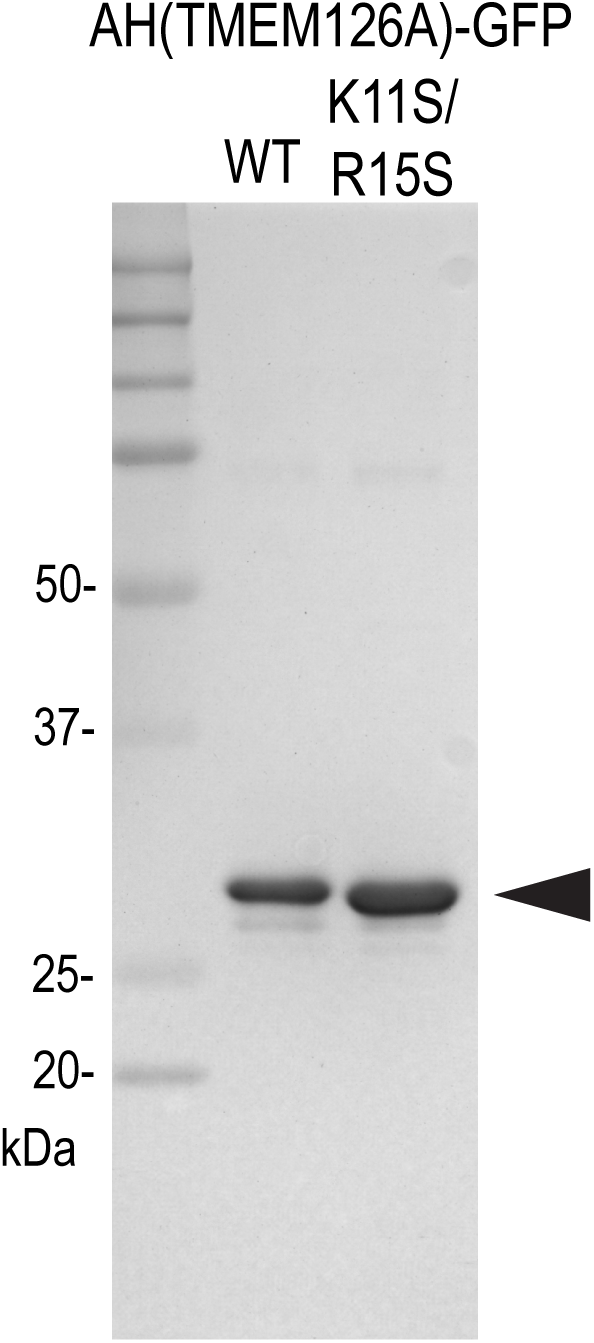
Purification of AH-GFP. Coomassie staining of the indicated AH-GFP recombinant proteins expressed and purified from bacterial cells.

**Fig. S11.**
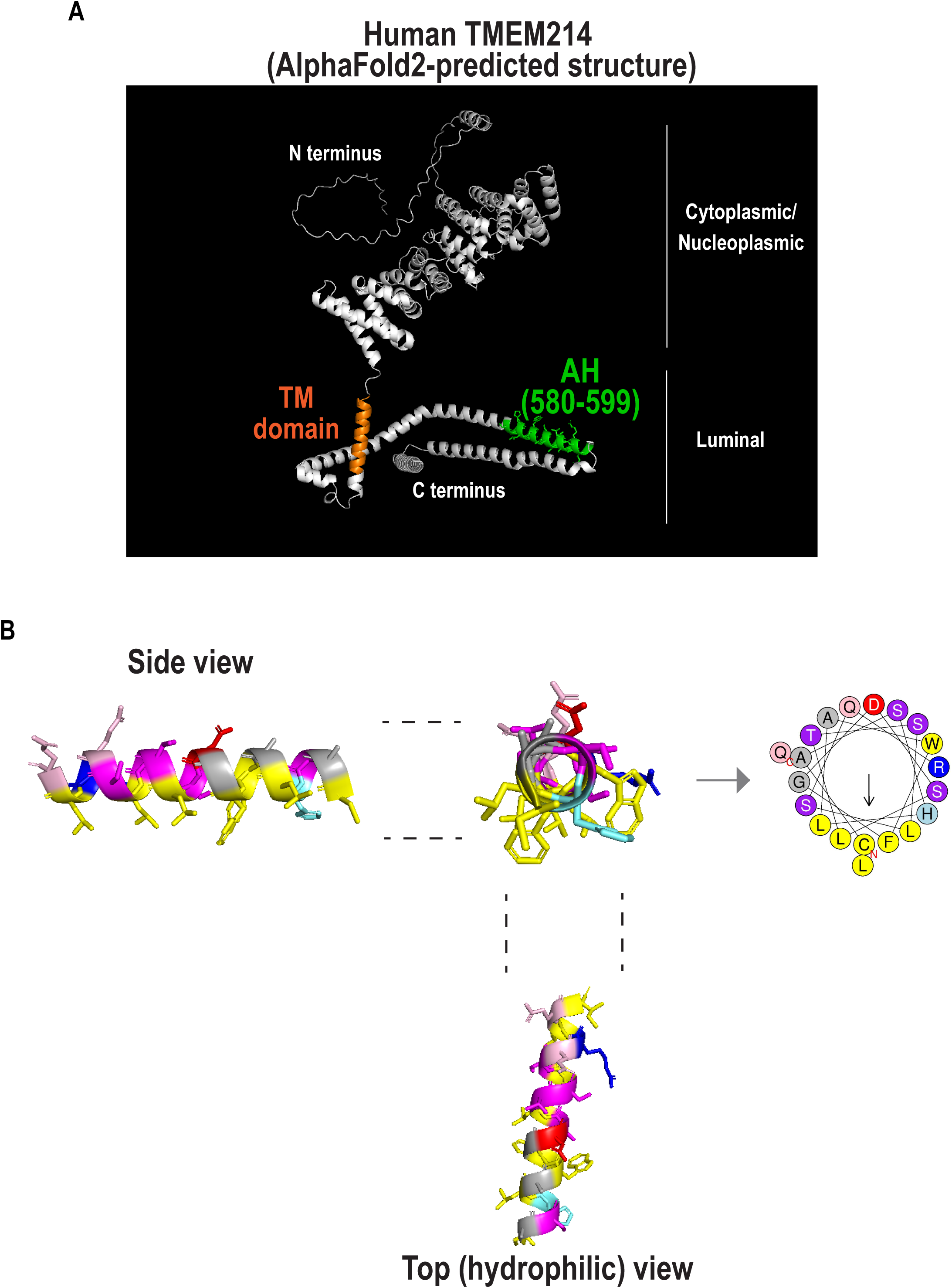
Putative AH in TMEM214 protein. (A) The putative AH is highlighted in the 3D structure of TMEM214 protein, predicted by AlphaFold2 and shown by PyMol. A putative transmembrane (TM) domain predicted by TMHMM is shown in brown. The N and C termini are predicted to face cytoplasm/nucleoplasm and NE/ER lumen, respectively, according to InterPro (ID: Q6NUQ4). (B) PyMol presentation of the TMEM214 AH. Amino acid residues are color-coded according to the wheel projection generated by HeliQuest.

**Fig. S12.**
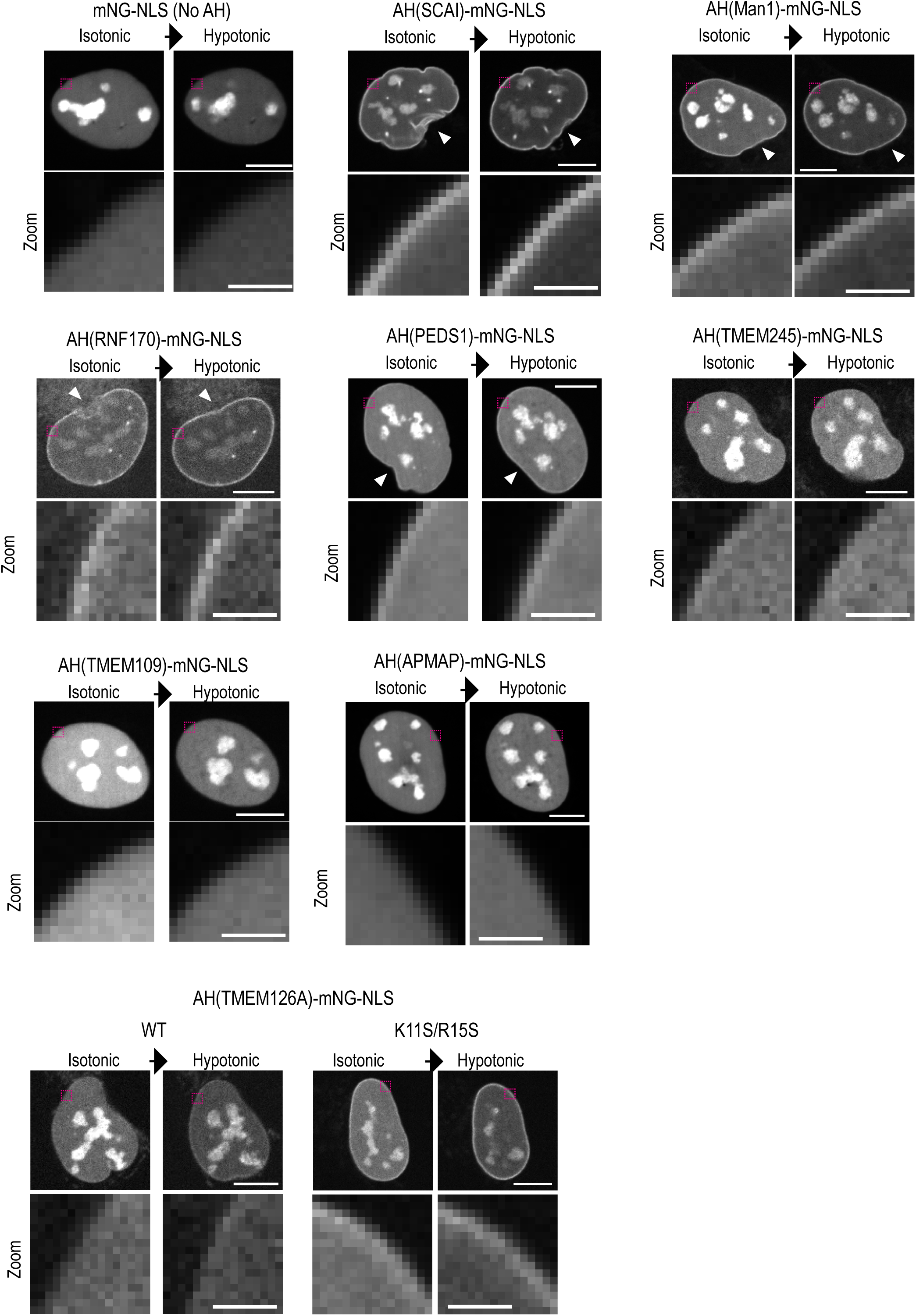
Hypotonic shock promotes AH binding to the inner nuclear membrane. Single z-section spinning disk confocal images of living U2OS cells expressing with the indicated AH-mNG-NLS constructs before and after the hypotonic shock. An arrowhead indicates the deformation of the nuclear rim wrinkle that is no longer present after hypotonic shock. Scale bars, 10 µm or 2 µm (zoom).

**Fig. S13.**
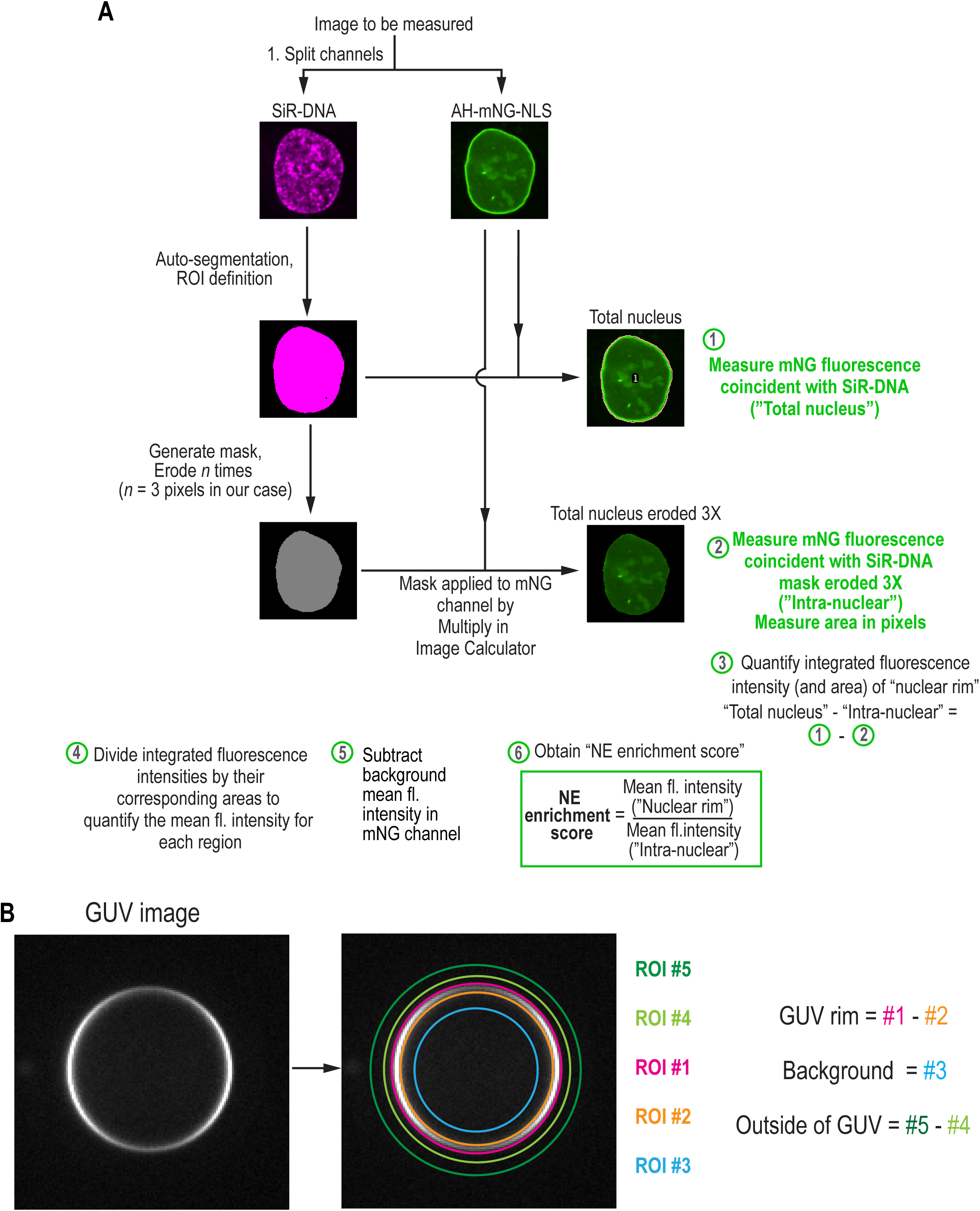
Flow chart of the NE enrichment score quantification and schematic of ROIs in AH-GUV binding quantification. (A) Flow chart of the NE enrichment score quantification. (B) Schematic illustration of ROIs in AH-GUV binding quantification. See methods for more detail.

